# A Cytoskeletal Vortex Drives Phage Nucleus Rotation During Jumbo Phage Replication in *E. coli*

**DOI:** 10.1101/2021.10.25.465362

**Authors:** Erica A. Birkholz, Thomas G. Laughlin, Sergey Suslov, Emily Armbruster, Jina Lee, Johannes Wittmann, Kevin D. Corbett, Elizabeth Villa, Joe Pogliano

**Affiliations:** Division of Biological Sciences, University of California, San Diego, La Jolla, CA; Leibniz Institute DSMZ–German Collection of Microorganisms and Cell Cultures, Braunschweig, Germany; Department of Cellular and Molecular Medicine, University of California, San Diego, La Jolla CA; Department of Chemistry and Biochemistry, University of California, San Diego, La Jolla CA; Howard Hughes Medical Institute, University of California, San Diego, La Jolla, CA

**Keywords:** bacteriophage, bacterial cell biology, bacterial cytoskeleton, phage replication

## Abstract

Vortex-like arrays of cytoskeletal filaments that drive cytoplasmic streaming and nucleus rotation have been identified in eukaryotes, but similar structures have not been described in prokaryotes. The only known example of a rotating intracellular body in prokaryotic cells occurs when nucleus-forming jumbo phages infect *Pseudomonas*. During infection, a bipolar spindle of PhuZ filaments drives intracellular rotation of the phage nucleus, a key aspect of the replication cycle. Here we show the *E. coli* jumbo phage Goslar assembles a phage nucleus surrounded by an array of PhuZ filaments resembling a vortex instead of a bipolar spindle. Expression of mutant PhuZ strongly reduces Goslar phage nucleus rotation, demonstrating that the PhuZ cytoskeletal vortex is necessary for rotating the phage nucleus. While vortex-like cytoskeletal arrays are important in eukaryotes, this work identifies the first known example of a coherent assembly of filaments into a vortex-like structure driving intracellular rotation within the prokaryotic cytoplasm.

## Introduction

Nucleus-forming *Pseudomonas* jumbo phages rely upon a bipolar spindle to position and rotate the phage nucleus that encloses their DNA and protects it from host defenses (Chaikeeratisak et al., 2021a). Analogous to a eukaryotic nucleus, the phage nucleus imparts strict separation of transcription from translation by enclosing phage DNA within a proteinaceous shell that excludes ribosomes (Chaikeeratisak et al., 2017a). This separation requires the export of mRNA to the cytoplasm for translation and the selective import of proteins required for transcription, DNA replication, and DNA repair. The phage nucleus grows as phage DNA is replicated inside and it is positioned in the center of the cell by a bipolar spindle composed of filaments of the tubulin-like PhuZ protein (Chaikeeratisak et al., 2017a; Erb et al., 2014; Kraemer et al., 2012). The phage nucleus protects the phage DNA by excluding restriction enzymes and DNA-targeting CRISPR-Cas in both *Pseudomonas* and *Serratia* jumbo phages (Malone et al., 2020; Mendoza et al., 2020; Nguyen et al., 2021). The phage nucleus and spindle are also important for jumbo phage diversification because they contribute to the evolution of new species through Subcellular Genetic Isolation and Virogenesis Incompatibility (Chaikeeratisak et al., 2021b).

Early in the nucleus-forming jumbo phage infection process, the phage nucleus is maneuvered towards the center of the host cell and oscillates in position once it reaches the midcell due to stochastic growth and shrinkage of PhuZ filaments in the bipolar spindle (Chaikeeratisak et al., 2017a; Erb et al., 2014; Kraemer et al., 2012). PhuZ, which forms a three-stranded polar filament with dynamic (+) and (-) ends, analogous to eukaryotic microtubules (Zehr et al., 2014), was first identified in phage 201φ2-1 infecting *Pseudomonas chlororaphis* and later shown to be conserved, along with the phage nucleus, in the related *Pseudomonas aeruginosa* phages ΦKZ and ΦPA3 (Chaikeeratisak et al., 2017b). The (-) ends of the spindle are positioned at each cell pole, while the (+) ends point toward the midcell. PhuZ filaments display dynamic instability during which the (+) ends rapidly depolymerize until returning to a growth phase, thereby allowing the spindle to position the phage nucleus in the center of the cell (Chaikeeratisak et al., 2017b; Erb et al., 2014; Kraemer et al., 2012).

During the later phases of the nucleus-forming jumbo phage replication cycle, PhuZ filaments exhibit treadmilling when they polymerize at the cell pole at a rate similar to depolymerization at the surface of the nucleus (Chaikeeratisak et al., 2019). Treadmilling filaments apply pushing forces to opposing sides of the phage nucleus, causing it to rotate (Chaikeeratisak et al., 2017a, 2019). Capsids, which assemble on the cell membrane, attach to treadmilling filaments of the PhuZ spindle and are trafficked to the surface of the phage nucleus where they dock for packaging of the genome (Chaikeeratisak et al., 2019). As the PhuZ spindle rotates the phage nucleus, capsids are distributed to different locations on its surface to promote efficient DNA packaging. These functions of the PhuZ spindle, including the remarkable capacity for driving intracellular rotation of the phage nucleus, are conserved in all three *Pseudomonas* jumbo phages described above and they are important for full efficiency of the replication cycle (Chaikeeratisak et al., 2017a, 2017b, 2019; Kraemer et al., 2012). To add to the intricacies of this replication mechanism, we recently described the discovery of phage bouquets (Chaikeeratisak et al., 2021c). These nearly spherical structures are formed by DNA-filled capsids with tails pointing inward, resembling a bouquet of flowers. Bouquets are established in the final stages of the nucleus-forming jumbo phage replication cycle when fully packaged capsids move from the surface of the phage nucleus to the regions adjacent to the nucleus. The interior of bouquets largely excludes ribosomes and cytoplasmic GFP (Chaikeeratisak et al., 2021c). Eventually, the cell lyses, releasing the progeny phages into the environment.

It is still unknown how widespread this replication pathway may be among jumbo phage; assembly of a phage nucleus has only been observed in the above-mentioned *Pseudomonas* phages and *Serratia* phage PCH45, while the PhuZ cytoskeleton has only been studied in *Pseudomonas*. In order to further understand if phage nucleus formation and positioning, bipolar PhuZ spindle assembly, intracellular rotation of the phage nucleus, capsid trafficking, and phage bouquets are conserved among more diverse jumbo phage, we sought out a nucleus-forming jumbo phage that infects *Escherichia coli*. Here we characterize the reproduction pathway of phage vB_EcoM_Goslar (Goslar) (Korf et al., 2019). We show that Goslar is a nucleus-forming jumbo phage that assembles a vortex-like cytoskeletal array instead of a bipolar spindle to drive phage nucleus rotation. PhuZ mutations that disrupt vortex assembly also disrupt phage nucleus rotation, linking the cytoskeletal vortex to a key process in the phage replication cycle. Our results show that the nucleus-forming phage infection pathway is likely widespread and diverse strategies have evolved to achieve intracellular rotation of the phage nucleus.

## Results

### E. coli jumbo phage Goslar forms a phage nucleus

*E. coli* phage vB_EcoM_Goslar (Goslar) was recently discovered and sequenced (Korf et al., 2019), revealing distantly related homologs of the PhuZ spindle protein and the major phage nucleus protein (Figures S1A and S1B). We studied the progression of Goslar through the lytic cycle in the *E. coli* lab strain MG1655 as well as its original host, APEC 2248 (APEC). We found host chromosomal DNA was largely degraded by 60 minutes post-infection (mpi) as a dense ball of phage DNA accumulated, visible by DIC as well as DAPI staining (Figures S1C-S1E), similar to infection of *Pseudomonas* by nucleus-forming phages (Erb et al., 2014; Kraemer et al., 2012). To determine whether the Goslar DNA ball is surrounded by a proteinaceous shell, we tagged a homolog of the major phage nucleus shell protein, Goslar gp189, (21.52% amino acid identity with gp105 of 201φ2-1; Figure S1A), with GFP on the N-terminus and expressed it from a plasmid by induction with 0.2 mM IPTG. In uninfected cells, fluorescence from the GFP-shell fusion was diffuse (Figure S2A). In infected cells imaged without added dyes, GFP-shell formed a distinct halo around the DNA density observed by DIC (Figures 1A and S2C). When visualized with FM4-64 and DAPI added to the imaging pad at 30, 60, and 90 mpi, GFP-shell surrounded the DNA (Figure 1D), suggesting that Goslar is a nucleus-forming phage.

**Figure 1.**
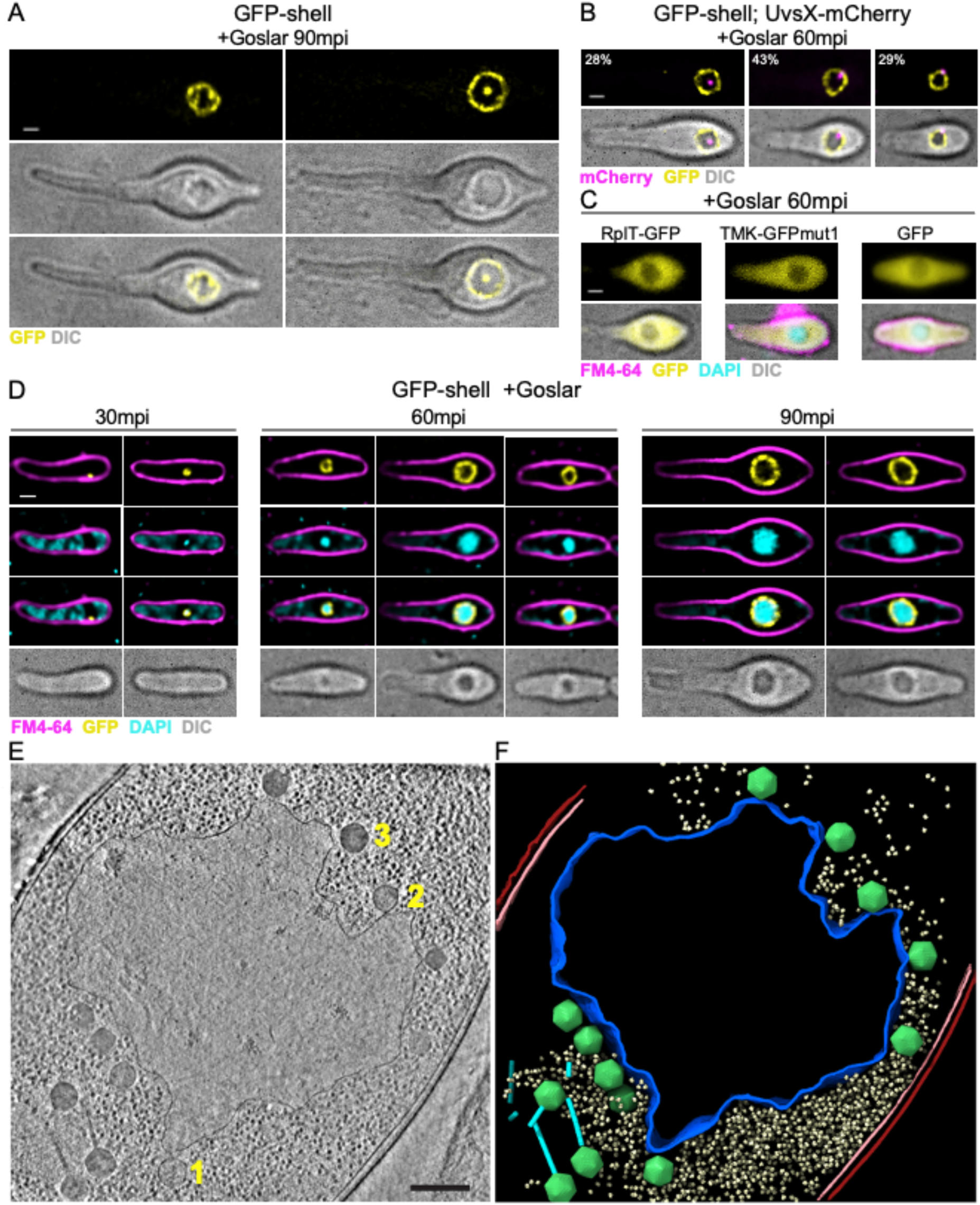
Goslar builds a phage nucleus separating DNA processes from translation and metabolism. (A) Goslar infecting *E. coli* (MG1655) expressing GFP-shell (gp189, yellow) at 0.2 mM IPTG and infected by Goslar for 90 minutes (mpi). White scale bars are 1 μm. (B) *E. coli* co-expressing GFP-shell and UvsX-mCherry at 0.2mM IPTG, infected by Goslar for 60 minutes. Displayed in the upper left is the percentage of the infected population that has UvsX-mCherry localized clearly to the interior of the GFP-shell halo (left), localized to the interior of the shell but on the periphery (center), and localized to the shell (right) (n=100). 100% of the time, UvsX-mCherry is colocalized with the phage nucleus. (C) *E. coli* expressing GFP fusions (yellow) with 50S ribosomal protein L20 (RplT), thymidylate kinase (TMK), and soluble GFP, and infected with Goslar for 60 minutes before being dyed with FM4-64 (4 μg/ml, membrane, magenta) and DAPI (2 μg/ml, DNA, cyan). (D) *E. coli* expressing GFP-shell, infected by Goslar for 30, 60, or 90 minutes then dyed with FM4-64 and DAPI. (E) Slice through a deconvolved tomogram of a phage nucleus in a 90 mpi Goslar-infected APEC cell. Inset scale bar is 250 nm (1 - empty capsid, 2 - partially filled capsid, 3 - nearly full capsid). (F) Annotation of the tomogram shown in (E). Outer and inner host cell membranes are colored red and pink, respectively. The phage nucleus shell is colored blue. Goslar capsids and tails are colored green and cyan, respectively. Host 70S ribosomes are colored pale yellow.

We next tested whether the Goslar phage nucleus separates proteins responsible for transcription and DNA maintenance from ribosomes and metabolic enzymes in the cytoplasm. Using fluorescent protein fusions, we found that the putative Goslar-encoded DNA repair enzyme UvsX (gp193) localizes with the Goslar phage nucleus. We co-expressed GFP-shell with UvsX-mCherry and found that UvsX-mCherry puncta were either clearly inside the GFP-shell (28%), inside the shell but also associated with it (43%), or colocalized with the surface (29%) (n=100; Figure 1B). Both fusions were diffuse when co-expressed in uninfected cells (Figure S2B). We observed the same association of UvsX with the phage DNA when UvsX was tagged with other fluorescent proteins (GFP, GFPmut1, mEGFP) on either terminus (Figures S2D-S2E). These results demonstrate that in 100% (n=100) of infections, UvsX was associated with the phage nucleus and never elsewhere in the cytoplasm, showing that it is selectively localized with the phage DNA.

The phage nuclei of the *Pseudomonas* jumbo phages exclude ribosomes and metabolic proteins (Chaikeeratisak et al., 2017a) so we examined the localization of the *E. coli* 50S ribosomal protein L20 (RplT) and thymidylate kinase (TMK) fused to a GFP, as well as GFP alone as a generally soluble protein. RplT-GFP, TMK-GFPmut1, and GFP were all diffuse in uninfected cells (Figure S2F) and they were largely excluded from the phage nucleus during infection (Figure 1C), demonstrating that the phage nucleus forms a compartment that is distinct from the cell cytoplasm. These data suggest that the Goslar phage nucleus successfully segregates DNA from ribosomes and metabolic enzymes, thereby uncoupling transcription from translation.

In order to examine the ultrastructure and molecular organization of the phage nucleus, we performed cryo-focused ion beam milling coupled with cryo-electron tomography (cryo-FIB-ET) on Goslar-infected APEC cells at late stages of infections (∼90 mpi). Despite the distant sequence homology of the shell (Figure S1A), the ultrastructure of the Goslar phage nucleus is strikingly similar to phage nuclei observed in the previously characterized *Pseudomonas* jumbo phages. As observed for those phage nuclei, the perimeter of the Goslar nucleus appears to be composed of a single layer of protein ∼6 nm thick. In agreement with our fluorescence microscopy results, the Goslar nucleus separates phage DNA from the cytosol and is devoid of host ribosomes (Figures 1E and 1F). Capsids containing various amounts of DNA, based on the density within the capsids, were docked on the surface of the phage nucleus shell, representing various stages of Goslar DNA packaging.

### Goslar forms a vortex-like array of PhuZ filaments

Goslar encodes a divergent homolog (gp201) of the phage cytoskeletal protein PhuZ (26.96% amino acid identity with 201φ2-1 gp59; Figure S1B). To determine whether this divergent homolog forms a bipolar spindle that organizes phage replication similarly to the nucleus-forming *Pseudomonas* jumbo phages, we tagged the N-terminus of Goslar gp201 with GFP and visualized it with fluorescence microscopy *in vivo*. Since the polymerization of tubulins like PhuZ occurs spontaneously above a critical concentration of monomers, we chose a concentration of IPTG (0.2 mM) for induction of GFP-PhuZ that resulted in spontaneous filament formation in less than 0.5% of cells (Figures S3B and S3C), but strongly labeled PhuZ filaments in infected cells (Figures 2A and 2B). During infection by Goslar, GFP-PhuZ was incorporated into filaments that organized into a vortex-like array around the phage nucleus (Figures 2A, 2B, and 2D). The percentage of cells containing GFP-PhuZ filaments increased from 19% (n=75) at 30 mpi to 97% (n=114) at 60 mpi (Figure 2C). At 30 mpi, one or two long filaments (yellow) could be observed in some cells (Figure 2A). By 60 mpi, GFP-PhuZ filaments were arranged into a vortex wrapping around the phage nucleus (visualized by DAPI-staining in cyan and as a DIC density) and terminating at the membrane with some reaching the cell poles (Figure 2A). This vortex-like cytoskeletal structure remained assembled at both 90 mpi and 120 mpi (Figure 2B). When GFP-PhuZ was expressed at lower induction levels, including 20 μM IPTG and 100 µM IPTG, the cytoskeletal vortex was still observed (Figure S3A). Even without any IPTG present, GFP signal from the fusion protein was most concentrated around the periphery of the phage DNA visualized by DIC, although the signal was very low. These results suggest that GFP-PhuZ filaments wrap around the phage nucleus and create a vortex-like cytoskeletal structure (Figure 2D) and that the vortex is not due to an artifact of overexpression.

**Figure 2.**
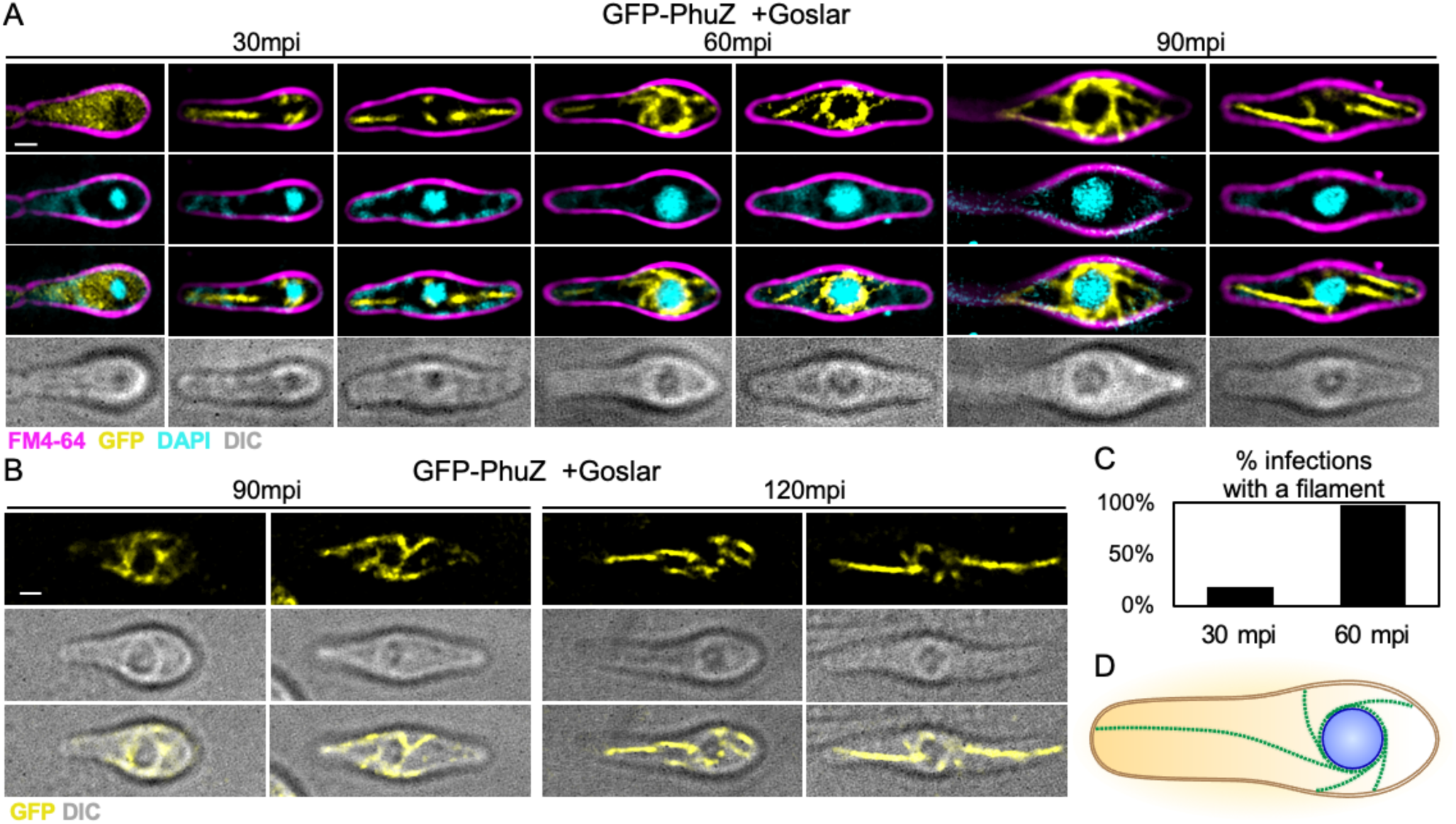
Goslar PhuZ forms a vortex-like cytoskeletal array. (A) *E. coli* (MG1655) expressing GFP-PhuZ at 0.2 mM IPTG and infected with Goslar for 30, 60, or 90 minutes prior to being dyed with FM4-64 and DAPI for 5-10 minutes. All scale bars are 1 μm. (B) *E. coli* (MG1655) expressing GFP-PhuZ and infected by Goslar for 90 and 120 minutes. (C) Percentage of 30 mpi (n=75) or 60 mpi (n=114) cells with a GFP-PhuZ filament over 0.3 μm. (D) Model of PhuZ cytoskeletal vortex. PhuZ filaments (green) extend radially from the phage nucleus (blue) to the cell membrane (gold).

To better visualize the spatial relationship between the PhuZ vortex and the phage nucleus, we simultaneously visualized the Goslar shell and PhuZ filaments by expressing GFP-shell and mCherry-PhuZ from the same plasmid with 0.2mM IPTG. At 90 mpi, the mCherry-PhuZ filaments wrapped around the GFP-shell and protruded towards the membrane in all directions (Figure 3A). To determine whether the choice of fluorescent protein fusion affected the co-localization, we also imaged mCherry-shell with GFP-PhuZ and found a similar organization (Figure 3B). In time course experiments, we found that at 30 mpi the shell of the phage nucleus had grown to the point that it was clearly distinguishable as a compartment containing DAPI-stained DNA before PhuZ filaments assembled a visible vortex around it (Figure 3C). By 60 mpi and beyond, the filaments colocalized with the phage nucleus surface and assembled a cytoskeletal vortex. This contrasts with the bipolar spindle assembled by nucleus-forming *Pseudomonas* phages, where filaments emanate from each pole of the cell and extend toward the phage nucleus at the cell center, providing the pushing forces necessary for both positioning and rotation (Figure 5D). Thus, the vortex-like organization of PhuZ filaments during Goslar infections represents a novel type of cytoskeletal structure found in prokaryotes (Chaikeeratisak et al., 2017a, 2017b; Erb et al., 2014; Kraemer et al., 2012).

**Figure 3.**
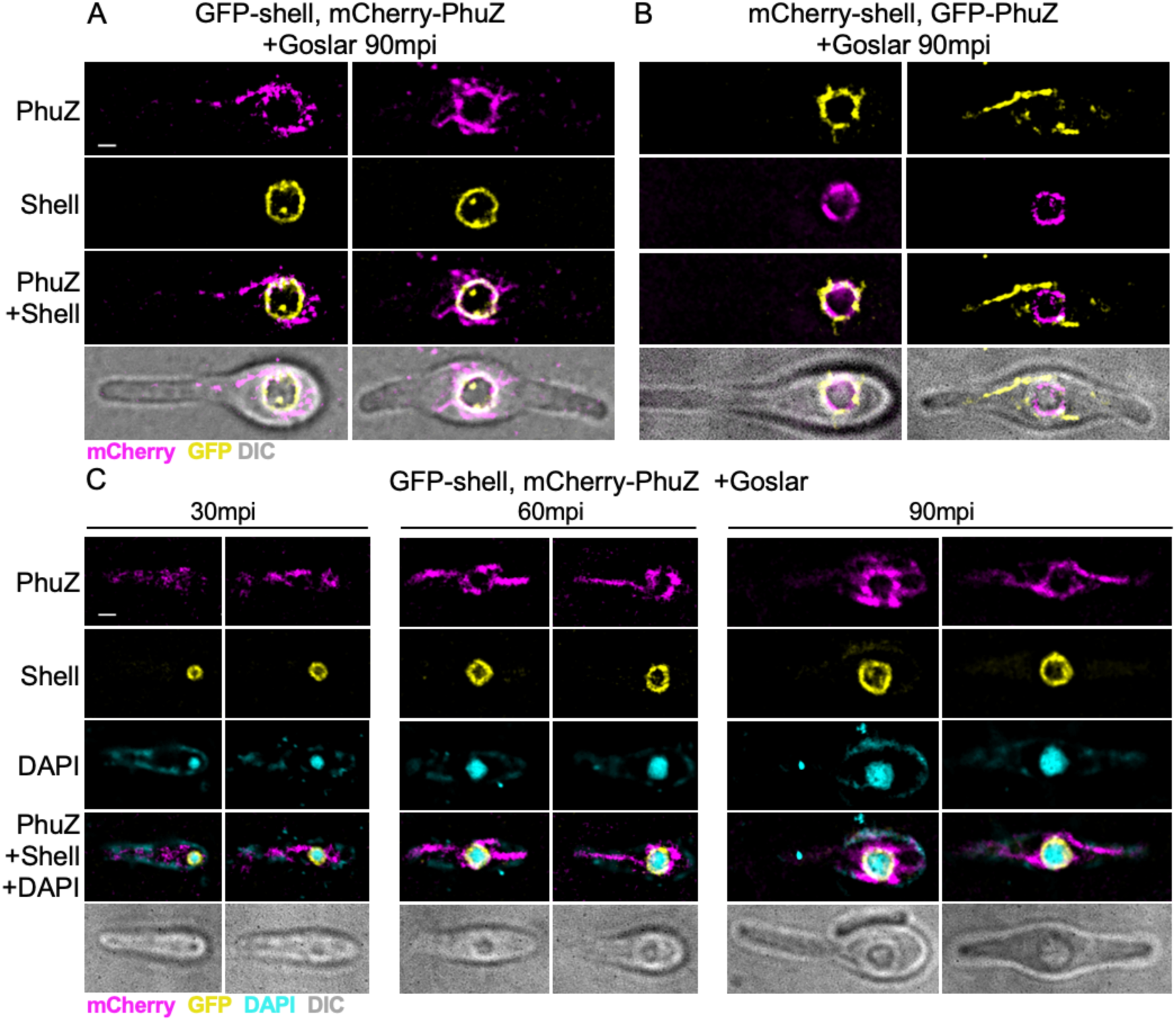
Colocalization experiments show the PhuZ cytoskeletal vortex wraps around the proteinaceous phage nucleus. (A) *E. coli* (MG1655) co-expressing GFP-shell (yellow) and mCherry-PhuZ (magenta) and infected with Goslar for 90 minutes. All scale bars are 1 μm. (B) *E. coli* co-expressing mCherry-shell and GFP-PhuZ and infected with Goslar for 90 minutes. (C) *E. coli* co-expressing GFP-shell (yellow) and mCherry-PhuZ (magenta) and infected with Goslar for 30, 60, or 90 minutes then dyed with DAPI (cyan).

### The Goslar nucleus is not positioned at midcell

The bipolar PhuZ spindle of *Pseudomonas* jumbo phages uses dynamic instability to center the phage nucleus. However, we measured the position of the Goslar phage nucleus and found that, unlike nucleus-forming *Pseudomonas* phages, the Goslar nucleus was not specifically positioned at any one location along the cell length, in neither MG1655 nor APEC (Figures 4A and 4C). In fact, the likelihood of finding a Goslar nucleus near midcell was not significantly greater than finding it elsewhere in the cell. Phage nuclei were excluded from the cell pole by at least 5% of the cell length (0.25 µm for a cell 5 µm long) based on the distance between the edge of the phage nucleus and the nearest cell pole (Figure 4E). The Goslar nucleus grew in size over time as DNA was replicated inside, ultimately reaching an average of ∼4 µm in diameter (Figures 4B and 4D). These data demonstrate that the Goslar-encoded distantly related PhuZ protein does not position the growing phage nucleus at midcell.

**Figure 4.**
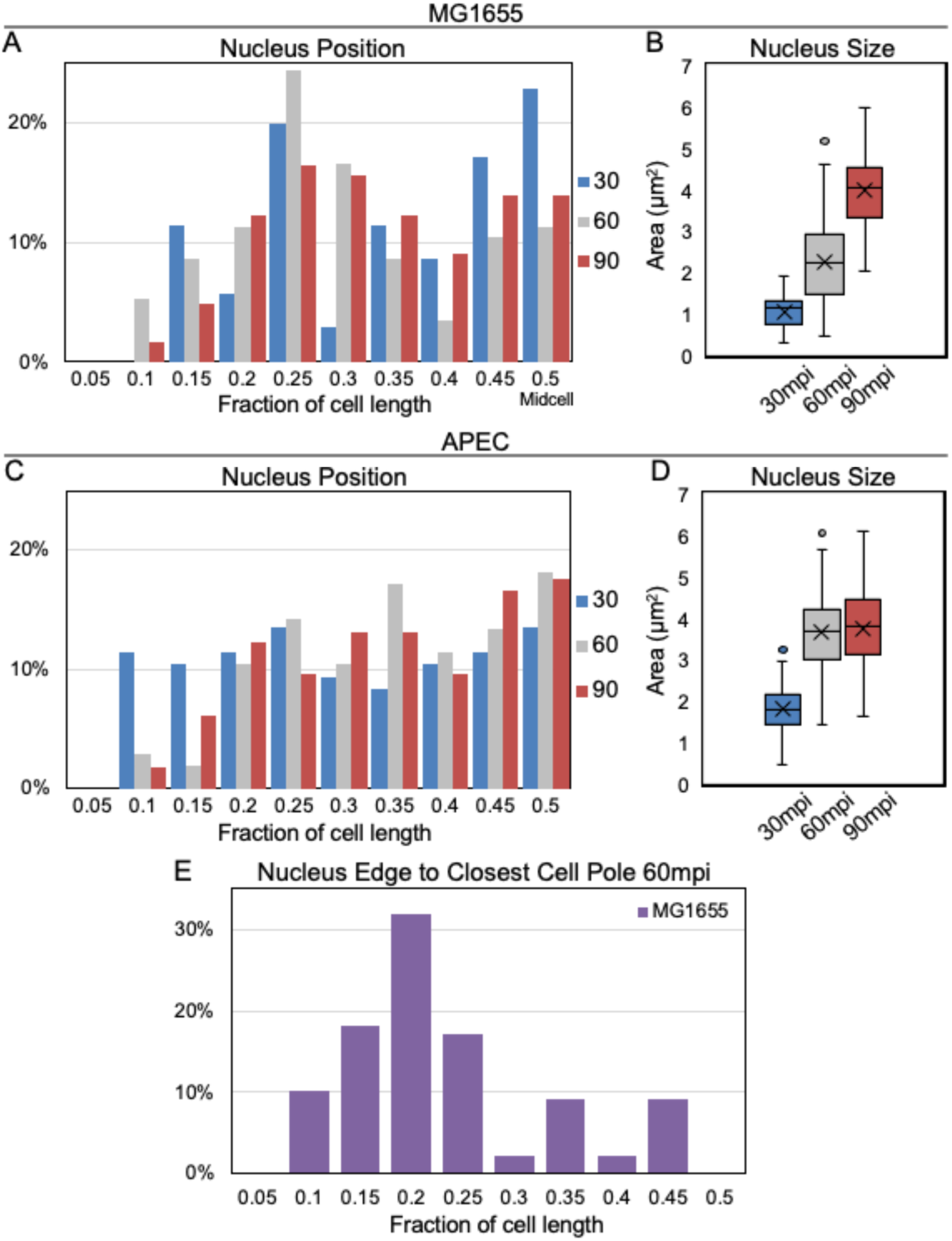
The Goslar nucleus is not positioned at midcell and is excluded from the cell pole. (A) Distribution of DIC phage nuclei positions along the lateral length of the cell (MG1655), in 0.05 μm bins. For each time point, there is no significantly greater chance of finding a phage nucleus near midcell (0.5) or quarter cell (0.25) than in the neighboring bins (30 mpi, n=35; 60 mpi, n=115; 90 mpi, n=122). (B) 2D area of the DAPI-stained phage nucleus in MG1655 at 30, 60, and 90 mpi (30 mpi; n=50, 60 mpi; n=114, 90 mpi; n=121). (C) Distribution of DIC phage nuclei positions along the lateral length of the APEC cell, in 0.05 μm bins. No significant enrichment occurs at midcell or any other bins (30 mpi; n=96, 60 mpi; n=105, 90 mpi; n=114). (D) 2D area of the DAPI-stained phage nucleus in APEC at 30, 60, and 90 mpi (30 mpi; n=115, 60 mpi; n=120, 90 mpi; n=145). (E) Distribution of distances between the edge of the phage nucleus to the closest cell pole, normalized to cell length. MG1655 infected for 60 minutes and visualized by DIC (n=88).

### Goslar nucleus rotation is dependent on vortex orientation and PhuZ function

Midway through the infection cycle, the phage nucleus of *Pseudomonas* phages 201φ2-1, ΦKZ, and ΦPA3 begins to rotate due to the opposing forces of the PhuZ spindle (Figure 5D) (Chaikeeratisak et al., 2017a, 2019). Rotation is coupled to the delivery of capsids to the surface of the phage nucleus, and disrupting PhuZ filament dynamics results in the production of fewer phage virions (Kraemer et al., 2012). Rotation is therefore an important and conserved aspect of the nucleus-forming jumbo phage replication cycle. We collected time-lapse images of the GFP-tagged shell every 4 seconds and found that the Goslar phage nucleus also rotates (Figure 5A, see Video 1). We also used DIC imaging to visualize and quantify nucleus rotation without the potential interference of any fluorescent stains or protein fusions (Figure 5B, see Video 2). The phage nucleus rotated with an average linear velocity of 50 ± 12 nm/s (measured at the nucleus periphery; n=20; Figure 5C). This rate of movement is nearly identical to the rates of nucleus rotation and PhuZ filament growth for the nucleus-forming *Pseudomonas* phages (Chaikeeratisak et al., 2019).

**Figure 5.**
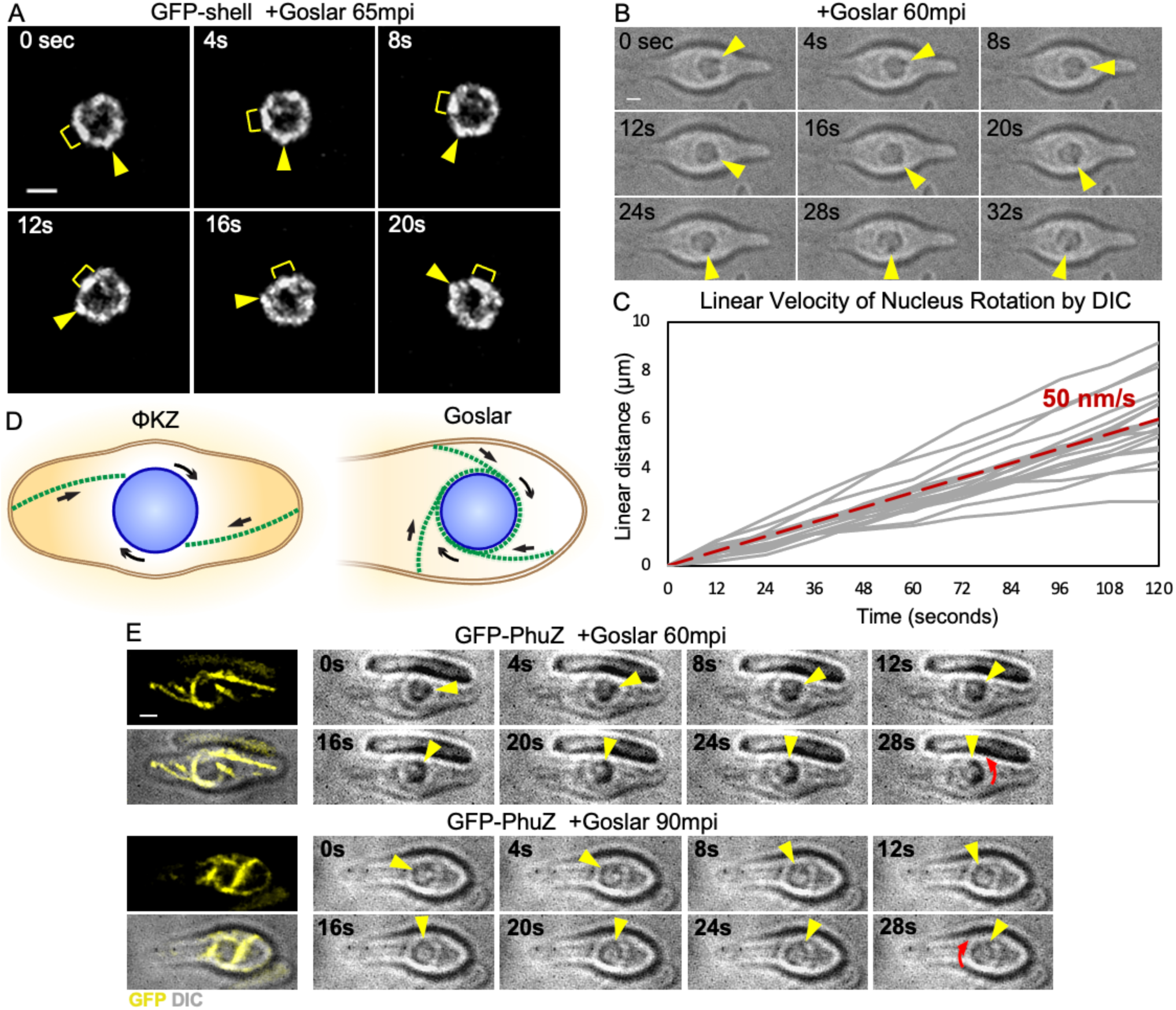
The Goslar phage nucleus rotates and the PhuZ vortex pushes against the cell membrane. (A) Time-lapse of the phage nucleus every 4 seconds for 20 seconds in *E. coli* (MG1655) expressing GFP-shell (white) infected with Goslar for 65 minutes. All scale bars are 1 μm. (B) DIC time-lapse every 4 seconds for 20 seconds in *E. coli* infected with Goslar for 60 minutes. (C) Linear velocity of nucleus rotation measured from DIC time-lapse, averaging 50 nm/s (n=20), red dotted line, individual measurements shown as gray lines. (D) Model of ΦKZ phage nucleus rotation by bipolar PhuZ spindle (left) and Goslar phage nucleus rotation by PhuZ cytoskeletal vortex (right). Arrows indicate the direction of forces applied to the phage nucleus. (E) GFP imaging coupled with DIC time-lapse every 4 seconds on *E. coli* expressing GFP-PhuZ; yellow arrowhead indicates DIC-dense spot to follow for rotation, red curved arrows in final panels indicate direction of rotation (CCW top cell, CW bottom cell).

We hypothesized a model of nucleus rotation for Goslar in which the PhuZ cytoskeletal vortex provides pushing forces tangentially to the surface of the phage nucleus (Figure 5D). This model predicts that the direction of nucleus rotation should be correlated with the orientation of filaments within the vortex, and that mutations that disrupt PhuZ filament dynamics by inactivating GTP hydrolysis will reduce the rate of rotation. To test the first prediction, we examined the direction of nucleus rotation relative to the orientation of the filaments. Nucleus rotation was observed using DIC time-lapse at 60 mpi and 90 mpi in cells induced with 0.2 mM IPTG to express GFP-PhuZ, and the PhuZ cytoskeleton was observed at a single time point just prior to the time-lapse. As shown in Figure 5E, the nucleus in the top panels rotated counterclockwise while the nucleus in the bottom panels rotated clockwise, and in both cases the cytoskeletal filaments were arranged in such a way that filament elongation would drive rotation of the phage nucleus in the corresponding direction. These results are consistent with a model in which the vortex of filaments drives rotation of the nucleus by pushing against the cell membrane (Figure 5D).

In order to further test this model, we mutated a conserved PhuZ aspartic acid residue, D202, to an alanine to disrupt the putative site of GTP hydrolysis. Expression of this mutant protein PhuZ(D202A) in uninfected cells yielded filaments in 56% of cells during induction with only 0.02 mM IPTG (Figures S4A and S4B), whereas filaments are only observed for wild-type PhuZ above 0.3 mM IPTG (Figure S3C). The ability of the mutant to assemble visible filaments when expressed at a much lower level compared to wild-type PhuZ suggests that the D202A mutation disrupts PhuZ polymer dynamics by inhibiting GTP hydrolysis. Based on our prior studies creating similar mutations in phage and plasmid-encoded tubulins (Erb et al., 2014; Kraemer et al., 2012; Larsen et al., 2007), we expected GFP-PhuZ(D202A) to behave as a dominant-negative mutant that could co-assemble with wild-type PhuZ to form inactive polymers. During Goslar infection in the presence of GFP-PhuZ(D202A) induced with 0.2 mM IPTG, the mutant protein failed to form a vortex (Figure 6A), yet phage nuclei positioning and growth appeared to be unaffected, compared to GFP-PhuZ (Figures S5A-S5E). However, in time-lapse microscopy, the intracellular rotation of phage nuclei was greatly reduced (Figure 6B, see Videos 3 and 4). In the presence of wild-type GFP-PhuZ, 92% (n=265; Figure 6C) of nuclei rotated visibly by DIC with an average linear velocity of 44 ± 9 nm/s (n=20; Figures 6D and S4C-S4D, black dotted lines), whereas with the expression of GFP-PhuZ(D202A), only 25% (n=101; Figure 6C) of nuclei appeared to rotate and for those 25%, they had an average linear velocity of 18 ± 9 nm/s (n=20; Figure 6D; Figure S4D, magenta dotted line), significantly slower than the wild-type. As a control, nucleus rotation in the presence of wild-type GFP-PhuZ (92% rotated, 44 ± 9 nm/s) behaved similarly in the absence of any fusion proteins (97% rotated, n=105, Figure 6C; 50 ± 12 nm/s, n=20, Figure 6D; Figure S4C, red dotted line). The disruption of function by expression of PhuZ(D202A) is likely not complete in some cells due to the presence of wild-type PhuZ expressed by the phage. These results are consistent with a model in which the PhuZ vortex is required for Goslar nucleus rotation.

**Figure 6.**
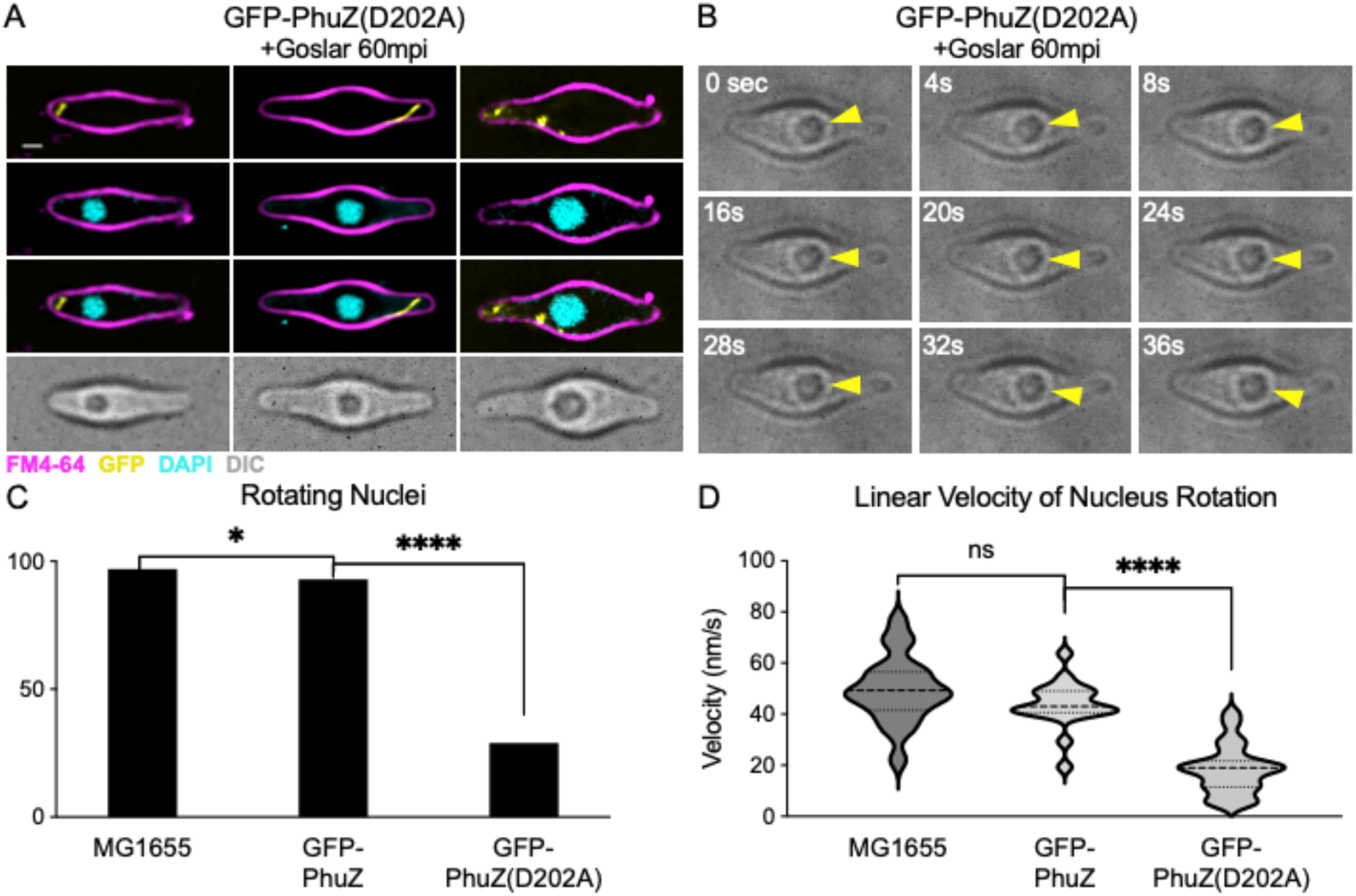
Mutant PhuZ(D202A) disrupts filament formation and nucleus rotation. *E. coli* expressing GFP-PhuZ(D202A) at 0.2mM IPTG and infected by Goslar for 60 minutes before being stained with FM4-64 and DAPI. All scale bars are 1 μm. (B) DIC time-lapse every 4 seconds for 36 seconds on *E. coli* expressing GFP-PhuZ(D202A) at 0.2 mM IPTG and infected by Goslar for 60 minutes. (C) Percentage of the infected cells at 60 mpi that had a rotating nucleus with any amount of progressive movement imaged by DIC time-lapse for *E. coli* with no plasmid (MG1655, n=105) or cells expressing GFP-PhuZ (n=164) or GFP-PhuZ(D202A) (n=101) (^*^ p value = 0.04, ^****^ p value <0.0001). (D) Linear velocity of the most progressively rotating nuclei at 60 mpi by DIC time-lapse for *E. coli* with no plasmid (MG1655, n=20) or cells expressing GFP-PhuZ (n=20) or GFP-PhuZ(D202A) (n=23). Violin plot generated and unpaired t tests performed using GraphPad Prism 9 (ns - p value >0.05, ^****^ p value <0.0001).

### Goslar forms bouquets of mature virions

As the PhuZ cytoskeleton of nucleus-forming *Pseudomonas* phages rotates the phage nucleus, it simultaneously delivers capsids to the surface of the nucleus for DNA packaging. Once filled with DNA, maturing phage particles assemble structures we recently termed “phage bouquets” (Chaikeeratisak et al., 2021c). We therefore examined if Goslar also forms phage bouquets late in the infection cycle. We tagged the putative Goslar capsid protein, gp41, at its C-terminus with GFP to determine whether Goslar virions also assemble bouquets. At 90 mpi, several stages of capsid organization were visible. In some cells, capsid-GFP was found around the nucleus, where the capsids were likely docked and in the process of packaging phage DNA (Figures 7A and S6A, left panels). We also observed the formation of spherical structures of capsid protein adjacent to the phage nucleus, or “bouquets”, with capsids also appearing within the centers of the bouquets (Figures 7A and S6A, middle panels, white arrowheads). Since *Pseudomonas* phage bouquets have not been shown to have capsids localized to the interior of the bouquet (Chaikeeratisak et al., 2021c), this could be a novel type of bouquet organization. Finally, in very swollen cells, capsids accumulated in the cytoplasm without any obvious organization (Figures 7A and S6A, right panels) To demonstrate whether capsids within these spherical structures are filled with DNA, we stained Goslar infections with 10 μg/ml DAPI at 90 mpi because the addition of DAPI prior to infection halted phage replication in MG1655 (Figure 7B). We observed a spherical pattern of faint DAPI staining, similar to the capsid-GFP localization and consistent with DNA-filled capsids arranged in a bouquet (white arrows). Curiously, no DAPI staining was visible in the interior of the bouquets, despite observing capsid-GFP localized in this region. We also visualized bouquet formation during Goslar infections of the APEC strain, which could be grown in 200 ng/ml DAPI without inhibiting phage replication. This resulted in very bright staining of spherically shaped DAPI-stained phage bouquets that also contained DAPI staining within the interior of the bouquet (Figure 7C), as predicted by capsid protein localization (Figure 7A). Cells with a single phage nucleus typically contained one or two bouquets that could be found on either one side of the phage nucleus or opposite sides of the phage nucleus (Figure 7C). The DAPI-staining of Figure 7C left panels is presented as a 3D reconstruction in Videos 5 and 6. An example of a cell with two phage nuclei and three phage bouquets is shown in Figure 7C with more examples in Figure S6B. The length and width of 105 bouquets at a median focal plane is displayed as a scatter plot (Figure S6C). Phage bouquets could become larger than the nucleus, reaching over 2 µm in width and over 5 µm in length, with a median value of 1.5 ± 0.75 µm (Figure S6D).

**Figure 7.**
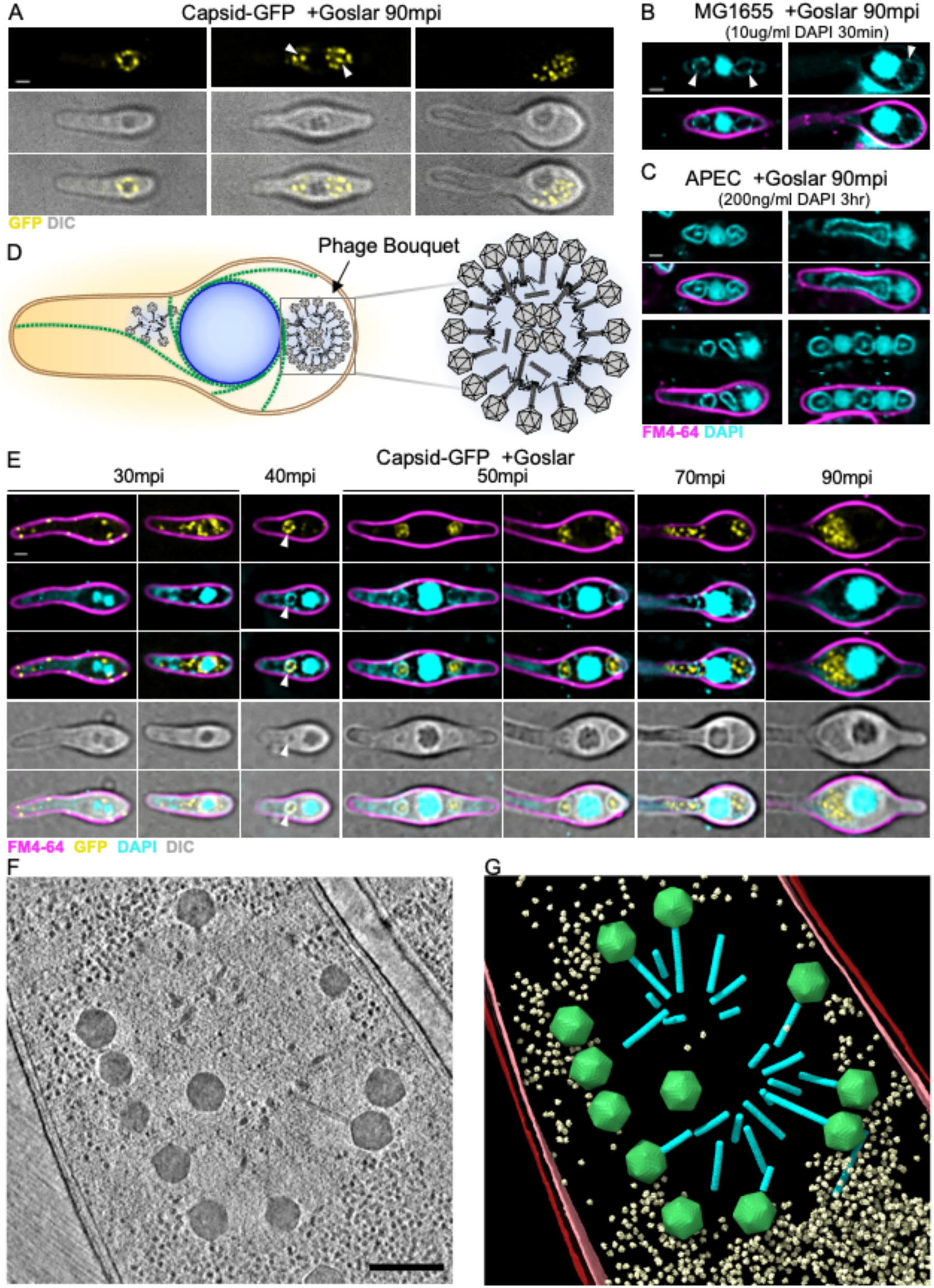
Goslar capsids migrate from the cytoplasm, surround the phage nucleus, and form phage bouquets. (A) *E. coli* (MG1655) expressing the putative capsid protein (gp41) fused to GFP (yellow) induced with 0.2 mM IPTG and infected with Goslar for 90 minutes. Three predominant localizations can be found; around the phage nucleus (left panels), in adjacent bouquets (middle panels), and filling more of the cytoplasm (right panels). White scale bars are 1 μm. (B) MG1655 infected with Goslar for 90 minutes then stained with 10 μg/ml DAPI for 30 minutes at room temperature. White arrowheads indicate faint bouquets. (C) APEC grown with 200 ng/ml DAPI for 90 minutes and then infected with Goslar for 90 minutes and imaged. Large, brightly fluorescent phage bouquets are formed. (D) Model of Goslar phage bouquet organization with tails packed together. (E) MG1655 expressing capsid-GFP (yellow) and infected with Goslar for 30, 40, 50, 60, 70, or 90 minutes before being stained with FM4-64 (magenta) and DAPI (cyan). (F) Slice through a deconvolved tomogram of a phage bouquet in a Goslar-infected APEC cell. Inset scale bar is 250 nm. (G) Annotation of the tomogram shown in (F). Outer and inner host cell membranes are colored red and pink, respectively. The phage nucleus shell is colored blue. Goslar capsids and tails are colored green and cyan, respectively. Host 70S ribosomes are colored pale yellow.

These observations suggest that Goslar capsids are organized into a sphere with the tails packed together as observed in the *Pseudomonas* phages (Chaikeeratisak et al., 2021c), but also containing capsids within the bouquet, not previously seen in the *Pseudomonas* phages (Figure 7D). To further investigate this model and better understand the stages of capsid localization, we simultaneously visualized capsid-GFP and DAPI staining at various times post infection (30, 40, 50, 70, and 90 minutes) (Figure 7E). We were not able to express capsid-GFP in APEC so we colocalized capsid-GFP with DAPI in MG1655. At early time points (30 mpi), the capsid fusion was devoid of DAPI staining and was found near the membrane and in the cytosol, or concentrated around the nucleus (Figure 7E). By 40 mpi, small circular bouquets stained with DAPI were found next to the DAPI-filled nucleus (Figure 7E, white arrow). Between 40 and 50 mpi, the small bouquets expanded and acquired internal capsids but only the exterior ring of capsids was stained with DAPI in MG1655. This time course resembles the trafficking of capsids from the membrane to the nuclear shell as described for the nucleus-forming *Pseudomonas* phages (Chaikeeratisak et al., 2019) and the assembly of “bouquets” of DNA-filled capsids in those phages (Chaikeeratisak et al., 2021c). Time-lapse microscopy of the capsid fusion without any dyes present confirmed the temporal progression of capsid localization from membrane or cytosol, to the periphery of the nucleus, and then to the adjacent bouquet(s) (Figure S6E, see Video 7).

Cryo-FIB-ET confirms that fully packaged capsids are arranged in a circular shape, forming bouquets (Figures 7F and 7G). Consistent with our fluorescence microscopy data, the interior of the observed bouquet is greatly depleted of host ribosomes compared to the surrounding cytosol. In the observed bouquet the majority of the tails point to the inside of the bouquet and the capsids to the exterior, but a capsid localized inside the bouquet is observed (Figures 7F and 7G). Given the sizes of the bouquets (>1.5 µm) compared to the thickness of FIB-milled lamellae (< 200 nm), we were unable to capture a bouquet in its entirety and ascertain its complete ultrastructure from our current cryo-FIB-ET dataset. Nevertheless, when combined with our fluorescence microscopy results, our cryo-FIB-ET data support the assembly of Goslar virions into double-layered bouquets at late stages of infection. Taken together, these data reveal that Goslar forms a distinctive type of prominent bouquet containing DNA-packed capsids within its center.

## Discussion

Nucleus-forming jumbo phages rely on a tubulin-based cytoskeleton for their complex subcellular organization. In *Pseudomonas* jumbo phages 201φ2-1, ΦPA3, and ΦKZ, PhuZ forms a dynamic bipolar spindle that positions the phage nucleus at midcell and delivers capsids to the shell of the phage nucleus while rotating it (Chaikeeratisak et al., 2017a, 2017b, 2019; Erb et al., 2014; Kraemer et al., 2012). Phage nucleus rotation occurs concomitantly with the delivery of newly assembled capsids to the surface of the nucleus (Chaikeeratisak et al., 2017a, 2019). We have proposed that this conserved feature of nucleus-forming jumbo phage replication allows capsids to be distributed uniformly around the nucleus surface for efficient DNA packaging into capsids. Intracellular rotation in prokaryotes has only been reported for *Pseudomonas* infected by nucleus-forming jumbo phages, where a bipolar array of PhuZ filaments provides the forces necessary for rotation. Here we characterize Goslar infecting *E. coli* and demonstrate that this phage also has a complex replication cycle with both a nucleus-like compartment and a tubulin-based cytoskeleton (Figure 8). Surprisingly, Goslar PhuZ filaments assemble a vortex-like cytoskeletal structure in which filaments wrap around the entire phage nucleus and project radially toward the membrane (Figure 8). The orientation of the cytoskeletal vortex is consistent with the hypothesis that it generates force against the membrane to cause nucleus rotation, and expression of the PhuZ(D202A) mutant impairs both vortex formation and nucleus rotation. Taken together, these results suggest that the function of the vortex-like cytoskeletal network of PhuZ filaments is to drive phage nucleus rotation. Previous work on phage 201φ2-1 has demonstrated that inactivating the PhuZ cytoskeleton eliminates both nucleus rotation and capsid trafficking thereby decreasing the number of phage progeny produced (Chaikeeratisak et al., 2019; Kraemer et al., 2012). In Goslar, the PhuZ cytoskeleton drives intracellular phage nucleus rotation, which is expected to facilitate capsid distribution. The PhuZ vortex may also participate in capsid trafficking, where radially projecting filaments would be ideal conduits for capsid migration from the cell membrane to the surface of the phage nucleus.

**Figure 8.**
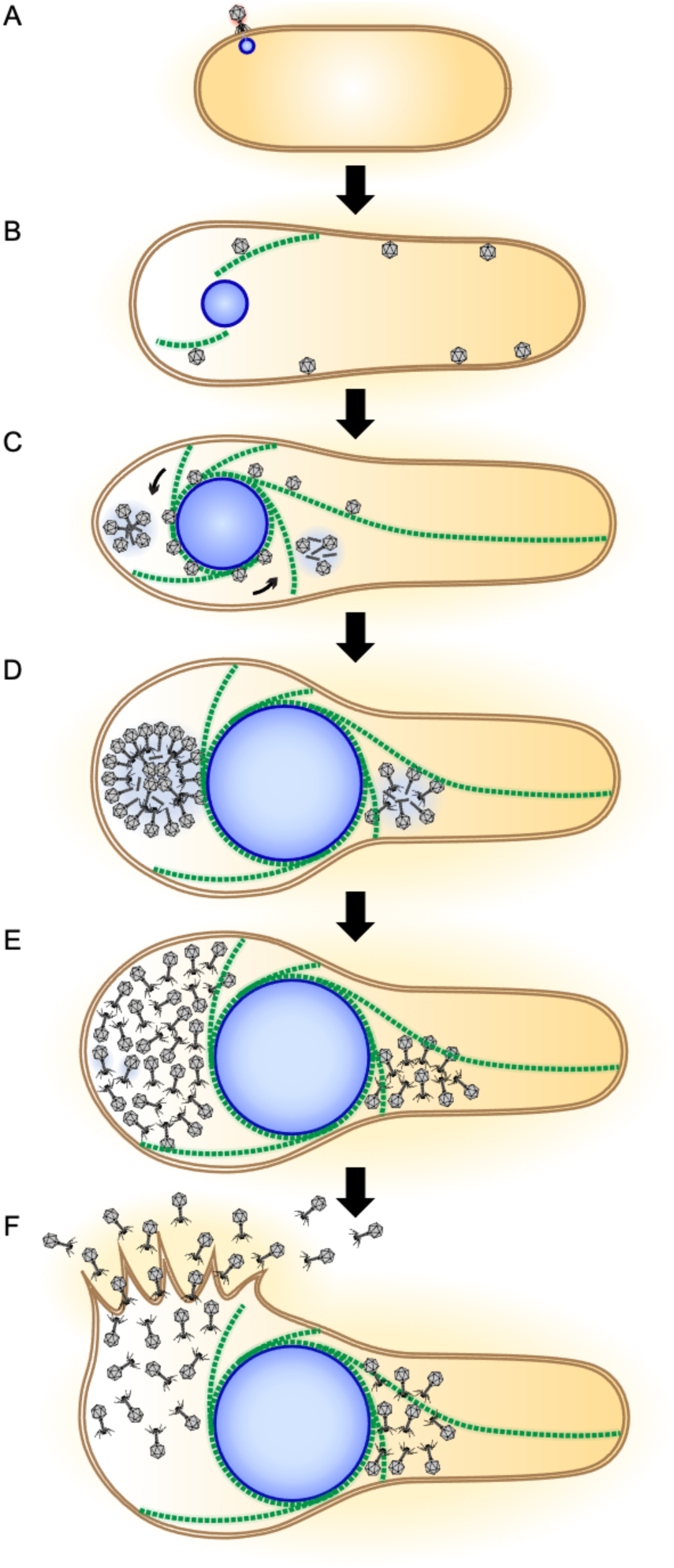
Model of the Goslar infection cycle. (A) The Goslar phage injects its DNA into an *E. coli* cell and the formation of a shell begins. (B) The shell grows in size as DNA replicates inside and the PhuZ vortex begins to form. Capsids form near the periphery of the cell and migrate towards the phage nucleus, possibly by trafficking along PhuZ filaments. (C) The PhuZ vortex is fully formed, wrapping around the phage nucleus. Capsids dock on the nuclear shell to be filled with DNA prior to localizing to the adjacent bouquets. (D) Large bouquets form with internally localized capsids on either one side or both sides of the phage nucleus. (E) Final assembly of the progeny virions is completed as they fill the cell in a more disordered fashion. (F) Lysis of the *E. coli* cell is achieved, releasing the progeny virions to find the next host.

A cytoskeletal vortex has not previously been described within any prokaryotes as far as we know. In eukaryotes, a vortex of cytoplasmic motion dependent on microtubules and actin has been shown to be critical for development by driving cytoplasmic streaming in the eggs of several species as well as in plant cells (Schroeder and Battaglia, 1985; Serbus et al., 2005; Stein et al., 2021; Woodhouse and Goldstein, 2013). The eukaryotic cytoskeleton is responsible for the rotation of the nucleus to align with cell polarity in motile cells, particularly fibroblasts (Fruleux and Hawkins, 2016; Gerashchenko et al., 2009; Kim et al., 2014; Levy and Holzbaur, 2008; Maninová et al., 2014; Wu et al., 2011) and has been observed in a vortex-like arrangement during misalignment (Kumar et al., 2014). A self-organizing vortex array of microtubules was demonstrated to spontaneously arise *in vitro* among purified microtubules with *Chlamydomonas* dynein c (Sumino et al., 2012) and in spatially confined droplets of *Xenopus* oocyte extract (Suzuki et al., 2017). The vortex of PhuZ filaments that organizes during Goslar infection is visually reminiscent of these *in vitro* microtubule vortex arrays.

While a cytoskeletal vortex is capable of providing intracellular rotation, it did not appear to play a role in phage nucleus positioning. The Goslar phage nucleus is not positioned at the midcell as it is for the nucleus-forming *Pseudomonas* phages. Instead, it is positioned along the entire lateral axis of the cell. Expression of the PhuZ(D202A) mutant, which significantly reduces nucleus rotation, does not affect nucleus positioning. Thus, the main function of the Goslar PhuZ cytoskeleton may be to provide the driving forces for nuclear rotation and perhaps capsid delivery, rather than midcell positioning.

At late stages of infection, Goslar virus particles accumulate in large spherical structures recently described for the nucleus-forming *Pseudomonas* phages and termed bouquets (Chaikeeratisak et al., 2021c). In phage bouquets, virions are arranged in a sphere with capsids on the outside and tails facing inward. Goslar bouquets appeared similar to ΦPA3 bouquets except Goslar’s contained capsids located in the center of the phage bouquets rather than only around the outside, suggesting that Goslar bouquets are organized with internal capsids oriented inversely relative to the larger outer layer, with tails packed together (Figures 8D and 7D). The role of phage bouquets is currently unknown, and the fact that they are not always detected in phages ΦPA3 and ΦKZ, and are only rarely detected in phage 201φ2-1, suggests that they are not essential for phage replication (Chaikeeratisak et al., 2021c). However, the discovery of prominent phage bouquets in Goslar, which is distantly related to the nucleus-forming *Pseudomonas* jumbo phages, suggests that bouquets likely offer an advantage to the phage.

The detailed characterization of the replication cycle of Goslar has brought to light a cytoskeletal vortex that drives phage nucleus rotation which is a key process in the nucleus-forming jumbo phage replication cycle. This work also demonstrates that the replication mechanism involving a phage nucleus and spindle is widespread across diverse hosts and that it likely confers a selective advantage since divergent strategies for rotation of the phage nucleus have evolved.

## Materials and Methods

### Growth conditions and bacteriophage preparation

Bacterial strains used in this study are listed in Tables S1 and S2. *Escherichia coli* strains APEC 2248 (APEC) and MG1655 were grown on Luria-Bertani (LB) plates containing 10 g Bacto-Tryptone, 5 g NaCl, 5 g Bacto-yeast extract, 16 g agar in 1 L ddH_2_O and incubated at 37°C overnight. Liquid cultures were obtained by inoculation of LB broth with one colony of *E. coli* from an LB plate. Lysates for Goslar were obtained from Johannes Wittmann at the DSMZ and were amplified by adding 15 µl high titer phage lysate to 300 µl APEC at OD_600_ 0.5, incubating at 37°C for 30 minutes, then adding 300 µl LB broth, plating 200 µl of the suspension onto each of 3 LB plates and incubating at 37°C overnight. 15 ml of Phage Buffer (PB) containing 10 mM Tris (pH 7.5), 10 mM MgSO_4_, 68 mM NaCl, and 1 mM CaCl_2_ was chilled on ice before 5 ml was added to each plate and left to soak at room temperature. After 4 hours, 3 ml of PB was added to each plate and after 2 more hours, the buffer was drawn off into a single tube. The lysate was clarified by pelleting the bacteria at 3220 x g for 10 minutes. The supernatant was filtered through 0.45 µm pores by syringe. 5 drops of chloroform from a pasteur pipette were added to the 10 ml lysate and shaken by hand for 1 minute. The mixture was spun at 3220 x g for 5 minutes and the aqueous phase containing the phage was removed to a clean tube and stored at 4°C.

### Plasmid constructions and bacterial transformation

Fluorescent-tagged phage proteins were synthesized into pDSW206 by Genscript and delivered as lyophilized plasmid. The plasmids were hydrated at ∼0.2 g/L with Tris-EDTA buffer and diluted 1:10 with ddH_2_O. Electroporation competent DH5 and MG1655 cells were prepared by washing with 10% glycerol and stored at -80°C. 30-50 µl of competent cells was combined with 1 µl of diluted plasmid and electroporated with 1.8 kV then incubated at 30°C in SOC media for 30-60 minutes before plating on LB with 100 µg/ml ampicillin and incubating overnight at 37°C.

### Single cell infection assay

*E. coli* strains containing a plasmid with fluorescent-tagged phage protein(s) were grown on 1% agarose pads supplemented with 25% LB and the desired IPTG concentration to induce protein expression. Cells were obtained from overnight incubation at 37°C on an LB plate with 100 µg/ml ampicillin. A colony was resuspended in 25% LB to an OD_600_ of ∼0.35 then 8 µl was spotted on the imaging pad and spread with the bottom of an eppendorf tube. Wild-type *E. coli* was grown without ampicillin or IPTG. The imaging pad was then incubated at 37°C for 1.5 hours without a coverslip in a humidor. 6 µl of Goslar lysate was added to the agarose pads and spread as before, then further incubated at 37°C to allow phage infection to proceed. At the desired time point, the slide was placed at room temperature and spotted with 7 µl of dye mix (2 µg/ml DAPI, 4 µg/ml FM4-64, 25% LB). Once dry after ∼5 minutes, a coverslip was put on the agarose pad and fluorescent microscopy was initiated. Data of static images and time-lapse imaging were collected and processed as described below.

### Live cell static image and time-lapse fluorescence microscopy

The DeltaVision Elite Deconvolution microscope (Applied Precision, Issaquah, WA, USA) was used to visualize the live cells. For static images, the cells were imaged with 12-15 slices in the Z-axis at 0.15 µm increments. For long time-lapse, imaging pads were prepared and infected as above and 30 minutes after the addition of Goslar, pads were coverslipped without dyes. The environmental control unit surrounding the microscope warmed the imaging space to 35°C. Fields adequate for imaging were marked and time-lapse imaging began 10 minutes post infection, with Ultimate Focus utilized. For short time-lapse, infections proceeded at 37°C for the indicated time before being coverslipped and imaged at room temperature one field at a time using Ultimate Focus. Images were processed by the aggressive deconvolution algorithm in the DeltaVision SoftWoRx 6.5.2 Image Analysis Program. Further image analysis and processing was performed in FIJI version 2.1.0/1.53c (Schindelin et al., 2012). Figure images were adjusted and layered in Adobe Photoshop.

### Cryo-electron tomography sample preparation

Ten agarose pads for infection were prepared as above (1% agarose, 25% LB) and spotted with 10 µl of APEC cells at an OD_600_ of ∼0.35 then incubated at 37°C for 1.5 hours in a humidor. 10 µl of Goslar lysate from the DSMZ was added and spread on each pad then incubated for another 1.5 hours until being removed to room temperature for cell collection. Infected cells were removed from the pads by addition of 25 µl of 25% LB and gentle scraping with the bottom of an eppendorf tube. All 10 pads were collected into one tube and after a portion was aliquoted, the remainder was centrifuged at 6000 x g for 45 seconds, resuspended with 1/4 volume of the supernatant, and a portion of that was diluted 1:1 in supernatant. Samples were delivered for plunging 20-30 minutes after removal from 37°C which significantly slows infection progression.

### Cryo-focused ion beam milling and electron tomography

Infected cells were prepared as described above (single cell infection assay) and at approximately 90 minutes post infection, 4-7 µl of cells were deposited on R2/1 Cu 200 grids (Quantifoil) that had been glow-discharged for 1 min at 0.19 mbar and 20 mA in a PELCO easiGlow device shortly before use. Grids were mounted in a custom-built manual plunging device (Max Planck Institute of Biochemistry, Martinsried) and excess liquid was blotted with filter paper from the backside of the grid for 5-7 seconds prior to freezing in a 50:50 ethane:propane mixture (Airgas) cooled by liquid nitrogen.

Grids were mounted into modified Autogrids (TFS) compatible with cryo-focused ion beam milling. Samples were loaded into an Aquilos 2 cryo-focused ion beam/scanning electron microscope (TFS) and milled to yield lamellae following published procedures for bacterial samples (Lam and Villa, 2021).

Milled specimens were imaged with a Titan Krios G3 transmission electron microscope (TFS) operated at 300 kV and equipped with a K2 directed electron detector (Gatan) mounted post Quantum 968 LS imaging filter (Gatan). The microscope was operated in EFTEM mode with a slit-width of 20 eV and using a 70 µm objective aperture. Automated data acquisition was performed using SerialEM-v3.8b11 (2005) and all recorded images were collected using the K2 in counting mode.

Tilt-series were acquired at either 4.27 Å and 5.34 Å per pixel. For the higher magnification tilt-series, images were acquired over a nominal range of +/-60° in 2° steps following a dose-symmetric scheme (Hagen et al., 2017) with a per-tilt fluence of 2.6 e^-^·Å^-2^ and total of ∼ 160 e^-^·Å^-2^per tilt-series. Lower magnification tilt-series were acquired similarly, but using a 1.5° tilt-step and per-tilt fluence of 1.8 e^-^·Å^-2^. Target defocus values were set for between -5 and -6 µm.

### Image processing and analysis of cryo-electron tomography data

Tilt-movies were corrected for whole-frame motion using Warp-v1.09 (Tegunov and Cramer, 2019) and aligned via patch-tracking using Etomo (IMOD-v4.10.28) (Mastronarde and Held, 2017). Tilt-series CTF parameters were estimated and tomograms reconstructed with exposure-filtering and CTF-correction using Warp-v1.09. For general visualization and membrane segmentation, tomograms were reconstructed using Warp’s deconvolution filter applied at default settings and downsampled to 20Å and 25Å per pixel from the original 4.27Å and 5.34Å pixel sizes, respectively.

Segmentation of host cell membranes and the phage nucleus perimeter from representative tomograms was performed by first coarsely segmenting using TomoSegMemTV (Martinez-Sanchez et al., 2014) followed by manual patching with Amira-v6.7 (TFS). Ribosomes, capsids, and tails were segmented using subtomogram analysis. For ribosomes, approximately 200 particles were manually selected from the respective tomograms and used to generate an *ab initio* reference using Warp-v1.09 and Relion-v3.1.1 (Scheres, 2012) following conventional procedures (Bharat and Scheres, 2016; Tegunov and Cramer, 2019). The references were used for template-matching with Warp-v1.09 against their respective tomograms. Template-matching results were curated to remove obvious false-positives (e.g., picks outside cell boundaries and cell membranes, etc.). Curated picks were aligned and classified using Relion-v3.1.1 to remove additional false-positives and refine their positions in the tomogram. For capsids, all particles were manually picked. Reference-generation and alignment of capsids was performed while enforcing icosahedral symmetry with Relion-v3.1.1 (despite the capsids possessing C5 symmetry) in order to promote convergence from the low number of particles. For the phage tails, the start and end points along the filament axis were defined manually and used to generate over-sampled filament models in Dynamo-v1.1514 (Castaño-DÍez et al., 2012, 2017). An initial reference for the tail was generated using Dynamo-v1.1514 from two full-length tails with clearly defined polarity. The resulting reference displayed apparent C6 symmetry, which was enforced for the alignment of all tails from a given tomogram using Dynamo-v1.1514 and Relion-v3.1.1. All interconversions of metadata between Warp/Relion and Dynamo formats were performed using scripts from the dynamo2m-v0.2.2 package (Burt et al., 2021). Final averages were placed back in the reference-frame of their respective tomograms using *dynamo_table_place*. Figures of the segmentation models were prepared using ChimeraX-v1.2.1 (Pettersen et al., 2021).

### Construction of PhuZ and shell phylogenetic trees

PSI-BLAST was run querying ΦKZ PhuZ (gp039) and shell (gp054). All phage-encoded hits were pulled and only those with both a PhuZ and a shell were aligned by MUSCLE and used to construct the trees in MEGA X (10.2.6) (Kumar et al., 2018; Stecher et al., 2020). The evolutionary history was inferred using the Neighbor-Joining method and bootstrapped with 1000 replicates. Evolutionary distances were calculated using the Poisson correction method.

### Quantification and Statistical Analysis

To quantify filaments of GFP-PhuZ or GFP-PhuZ(D202A), non-deconvolved DeltaVision image files were opened in FIJI version 2.1.0/1.53c using the Bio-formats importer. Images were automatically scaled at 15.456 pixels per µm and filaments were measured in the GFP channel at the slice containing the filament. For the percentage of the infected population that had a rotating nucleus, an A/B test was performed and a p value < 0.05 was the threshold for statistical significance.

Measurement of the linear velocity of nucleus rotation was performed on DIC images taken every 4 seconds. Dark protrusions on the surface of the phage nucleus were traced by hand over 12 second intervals (3 images) using the segmented line tool and their lengths recorded. Total distance travelled by points on the nuclear surface was divided by total time (120 seconds) and averaged with standard deviation calculated. Data is displayed as a violin plot produced in GraphPad Prism 9.2.0 for Mac OS X. An unpaired t-test was performed to obtain p values (p < 0.05, significant).

## Author Contributions

E.A.B., T.G.L., S.S., E.A., and J.L. conducted experiments and analyzed data. E.A.B. and J.P. conceptualized the original manuscript. J.W. provided the phage for study. E.A.B., T.G.L., E.A., J.W., K.D.C., E.V., and J.P. contributed to editing the manuscript.

The authors declare no conflict of interest.

This article contains supporting information online.

## Acknowledgements

This work was supported by the National Institutes of Health R01-GM129245 (to JP and EV) and R21-AI148814 (to KDC) and the National Science Foundation MRI grant NSF DBI 1920374 (to EV). EV is an investigator of the Howard Hughes Medical Institute. We acknowledge the use of the UC San Diego cryo-EM facility, which was built and equipped with funds from UC San Diego and an initial gift from the Agouron Institute. We thank Arshad Desai, Justin R. Meyer, and Amy M. Prichard for their helpful suggestions and comments on the manuscript.

---

**Table S1.** Bacteria, viruses, and plasmids used in this study.

| Strain or plasmid | Source or reference | Identifier |
| --- | --- | --- |
| *E. coli* MG1655 | CGSC | CGSC 6300 |
| *E. coli* APEC 2248 (APEC) | DSMZ | DSM 103255 |
| 5-alpha competent *E. coli* | New England Biolabs | Cat#C2987H |
| vB_EcoM_Goslar | Johannes Wittmann (DSMZ) (Korf et al., 2019) | DSM 104658 |
| pDSW206 plasmid | David Weiss (Weiss et al., 1999) | N/A |

**Table S2.**
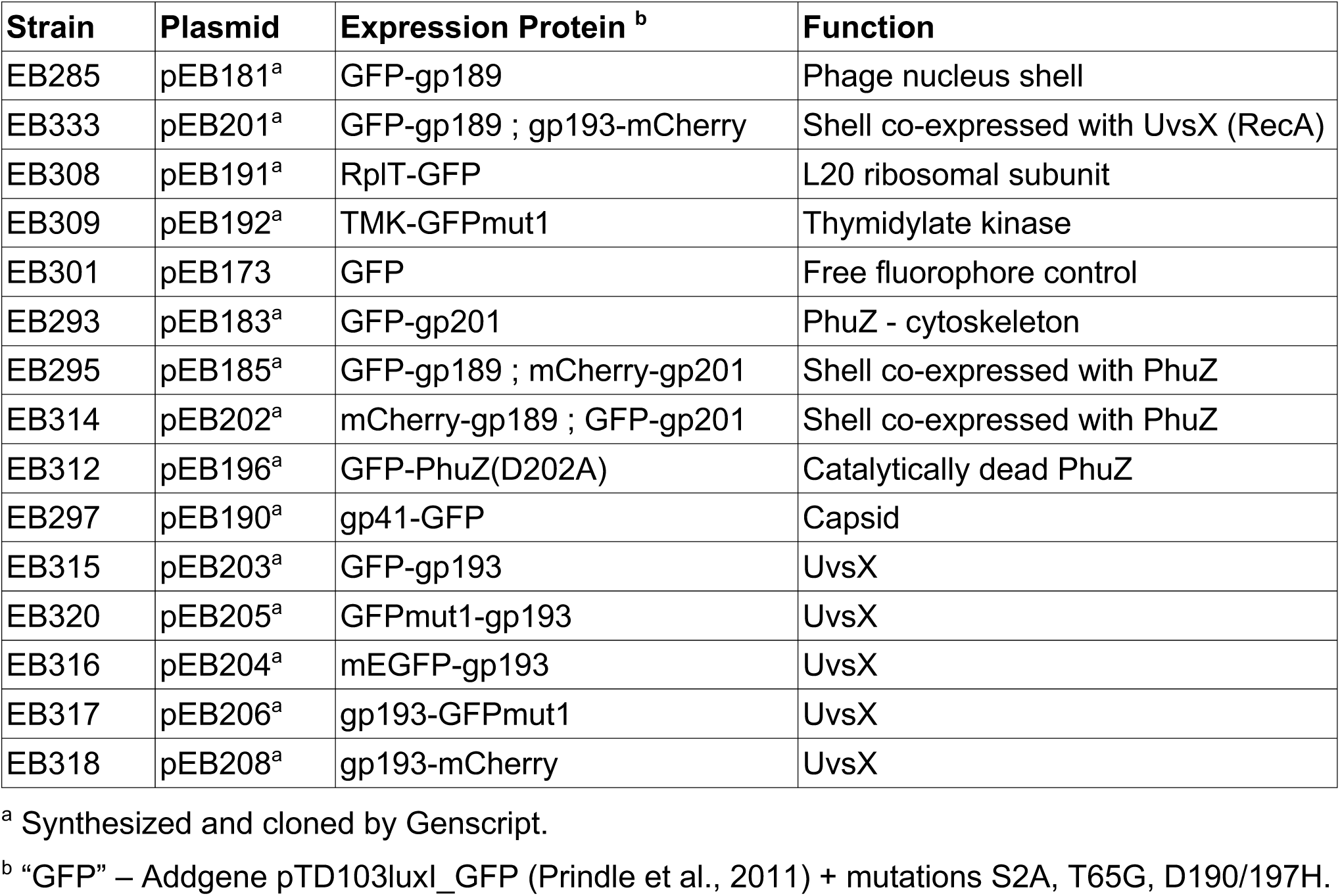
Plasmids generated for this study.

## Supplemental Figures

**Figure S1.**
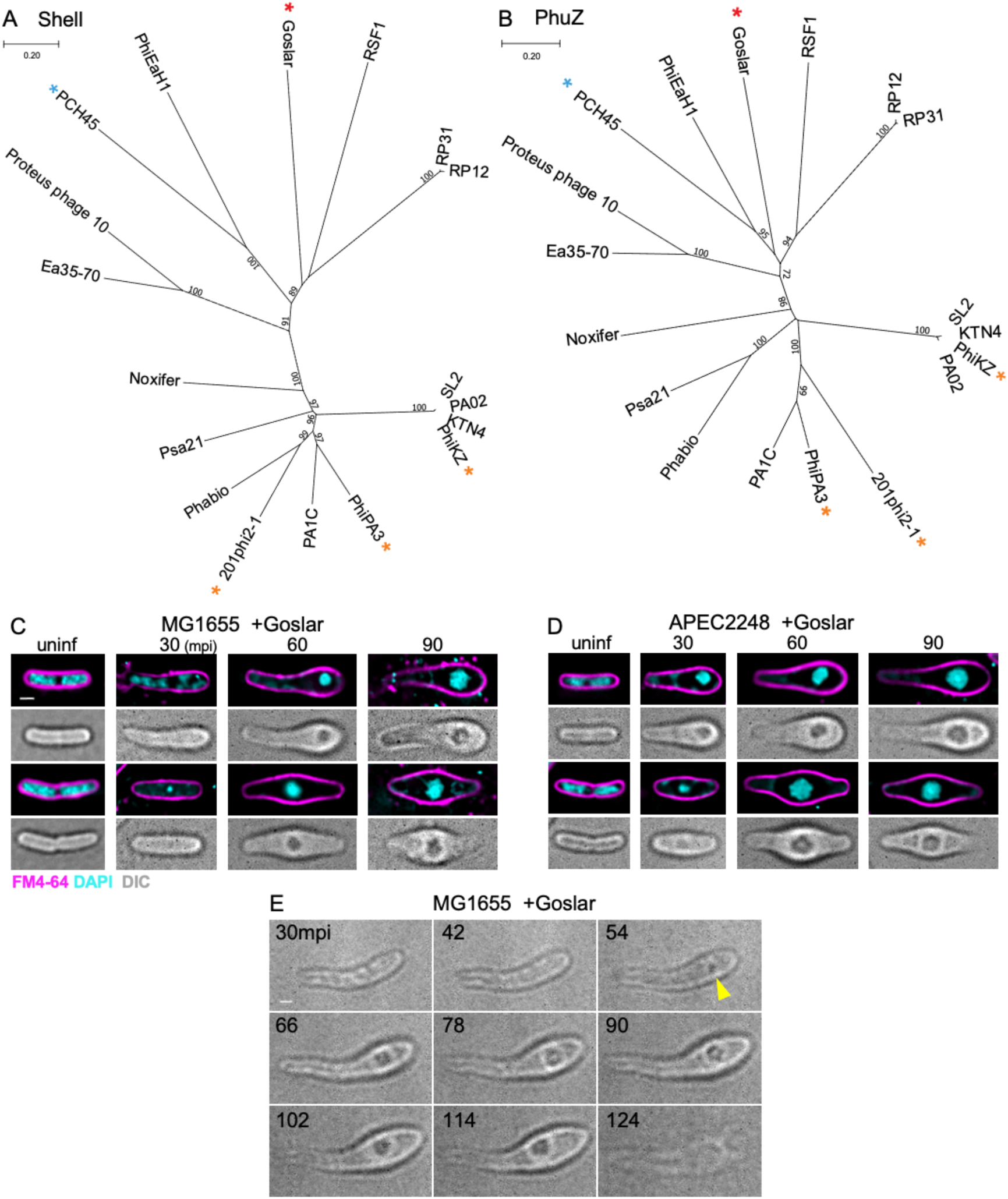
Goslar encodes distant homologs of the major phage nucleus shell protein and tubulin-like PhuZ protein and forms a dynamic DNA density during infection. (A & B) Unrooted phylogenetic trees of phage nucleus shell proteins (A) or phage tubulin PhuZ proteins (B) with bootstrap values (1000 replicates). Red asterisks: Goslar, gold asterisks: characterized nucleus-forming *Pseudomonas* phages, blue asterisks: nucleus-forming *Serratia* phage. (C & D) Two examples of MG1655 cells (C) and APEC cells (D) either uninfected (uninf) or infected by Goslar for 30, 60, or 90 minutes (mpi) then stained with FM4-64 (membrane, magenta) and DAPI (DNA, cyan). A concentrated mass of DNA appears only after addition of phage lysate and this mass can be observed without any fluorescent stains using DIC. All scale bars are 1 μm. Goslar replicates equally well in both strains. (E) DIC time-lapse every 12 minutes for 94 minutes of MG1655 infected with Goslar for 30 minutes prior to imaging. The DIC density shows that the nucleoid grows in size and moves around over time as the cell bulges.

**Figure S2.**
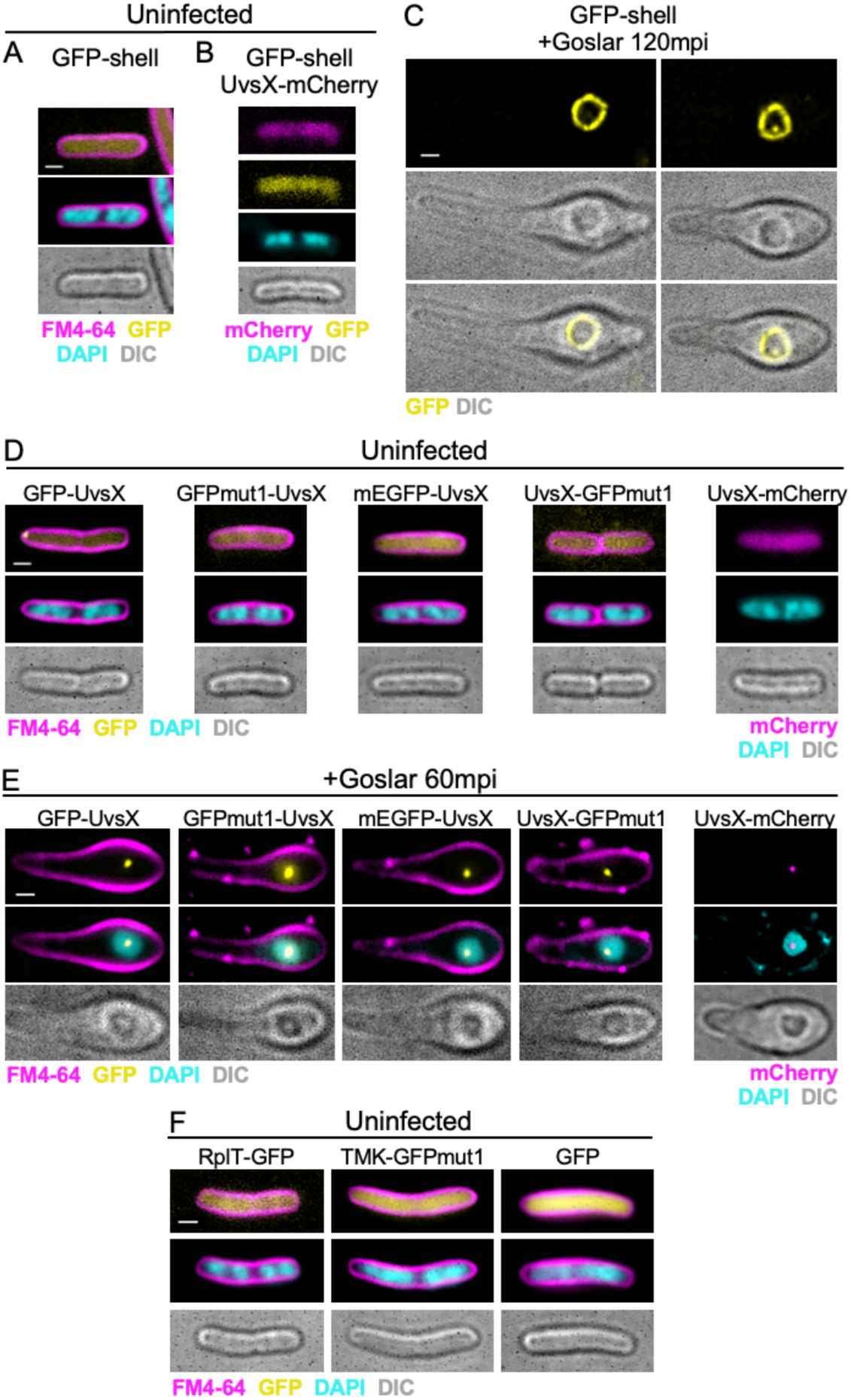
Control experiments show that the fusion proteins do not form specific structures in uninfected cells and that the choice of fusion does not alter the results. (A) *E. coli* (MG1655) expressing GFP-shell induced with 0.2 mM IPTG for 1.5 hours (uninfected). All scale bars are 1 μm. (B) *E. coli* co-expressing GFP-shell with UvsX-mCherry and induced at 0.2 mM IPTG for 1.5 hours then stained with DAPI (uninfected). (C) *E. coli* expressing GFP-shell induced with 0.2 mM IPTG and infected by Goslar for 120 minutes. (D) *E. coli* expressing each UvsX fusion indicated and induced at 0.2 mM IPTG for 1.5 hours then stained with FM4-64 and DAPI, or just DAPI for UvsX-mCherry (uninfected). Fusions to different fluorescent proteins (mCherry, mEGFP, GFPmut1) are generally uniformly distributed throughout the cell, only GFP-UvsX forms a polar punctum. (E) *E. coli* expressing the listed UvsX fusion protein at 0.2 mM IPTG, infected by Goslar for 60 minutes then stained with FM4-64 and DAPI. Fusing different fluorescent proteins (mCherry, GFP, mEGFP, GFPmut1) to UvsX at either the N- or C-terminus produces the same results. (F) *E. coli* expressing 50S ribosomal protein L20 (RplT-GFP), thymidylate kinase (TMK-GFP), or GFP alone, induced at 0.2 mM IPTG for 1.5 hours then stained with FM4-64 and DAPI. These fusions are generally uniformly distributed throughout the cell.

**Figure S3.**
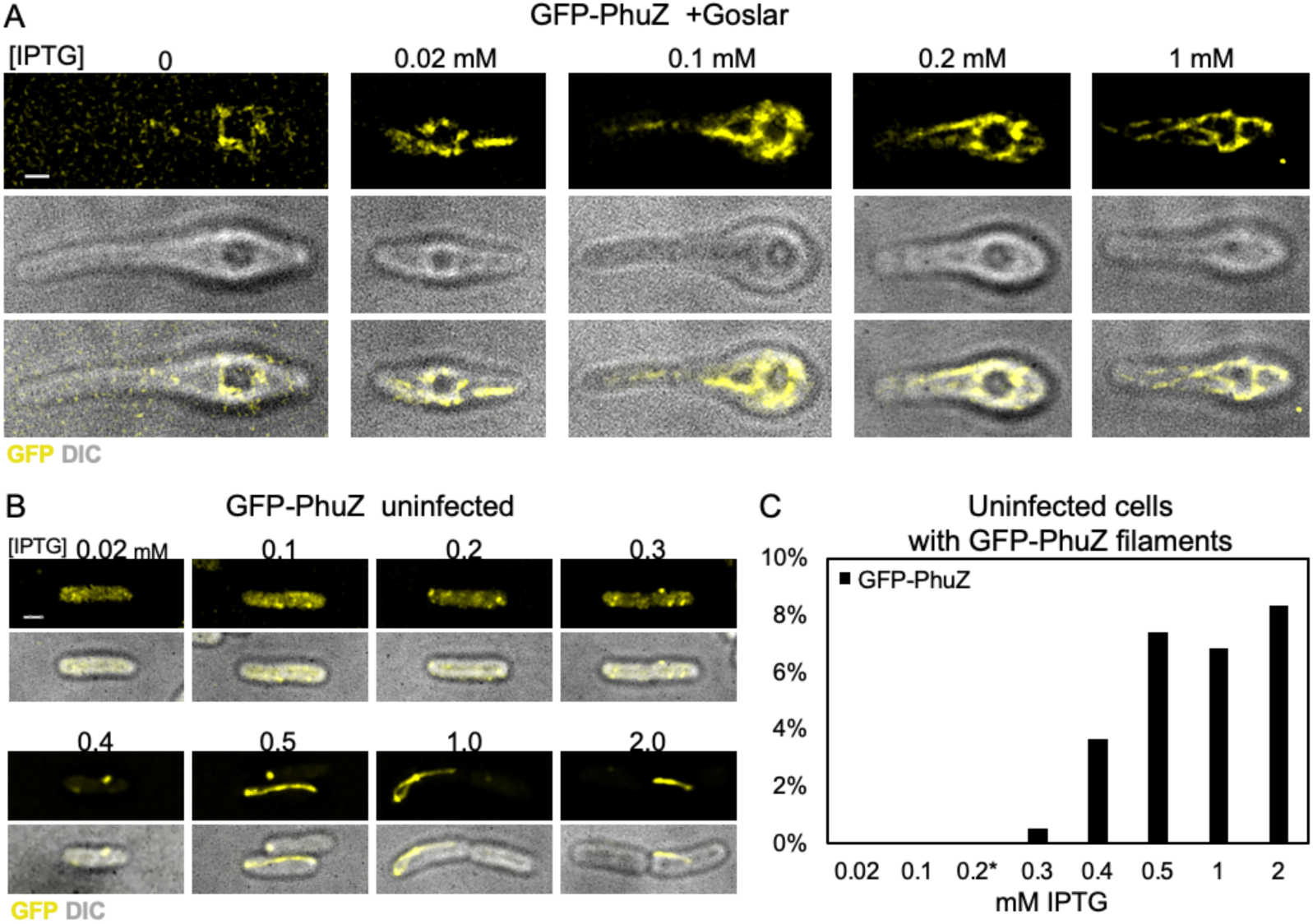
Assembly of GFP-PhuZ in uninfected and infected cells. (A) GFP-PhuZ accumulates around the phage nucleus even when expressed at very low levels. All levels of induction by IPTG show a similar vortex-like phenotype. All scale bars are 1 μm. (B) Uninfected *E. coli* (MG1655) expressing GFP-PhuZ (yellow) does not form filaments over 0.3 μm in length until 0.3 mM IPTG. (C) Percent of uninfected cells with a PhuZ filament over 0.3 μm at IPTG concentrations from 0.02 - 2.0 mM (0.02, n=78; others, n > 100).

**Figure S4.**
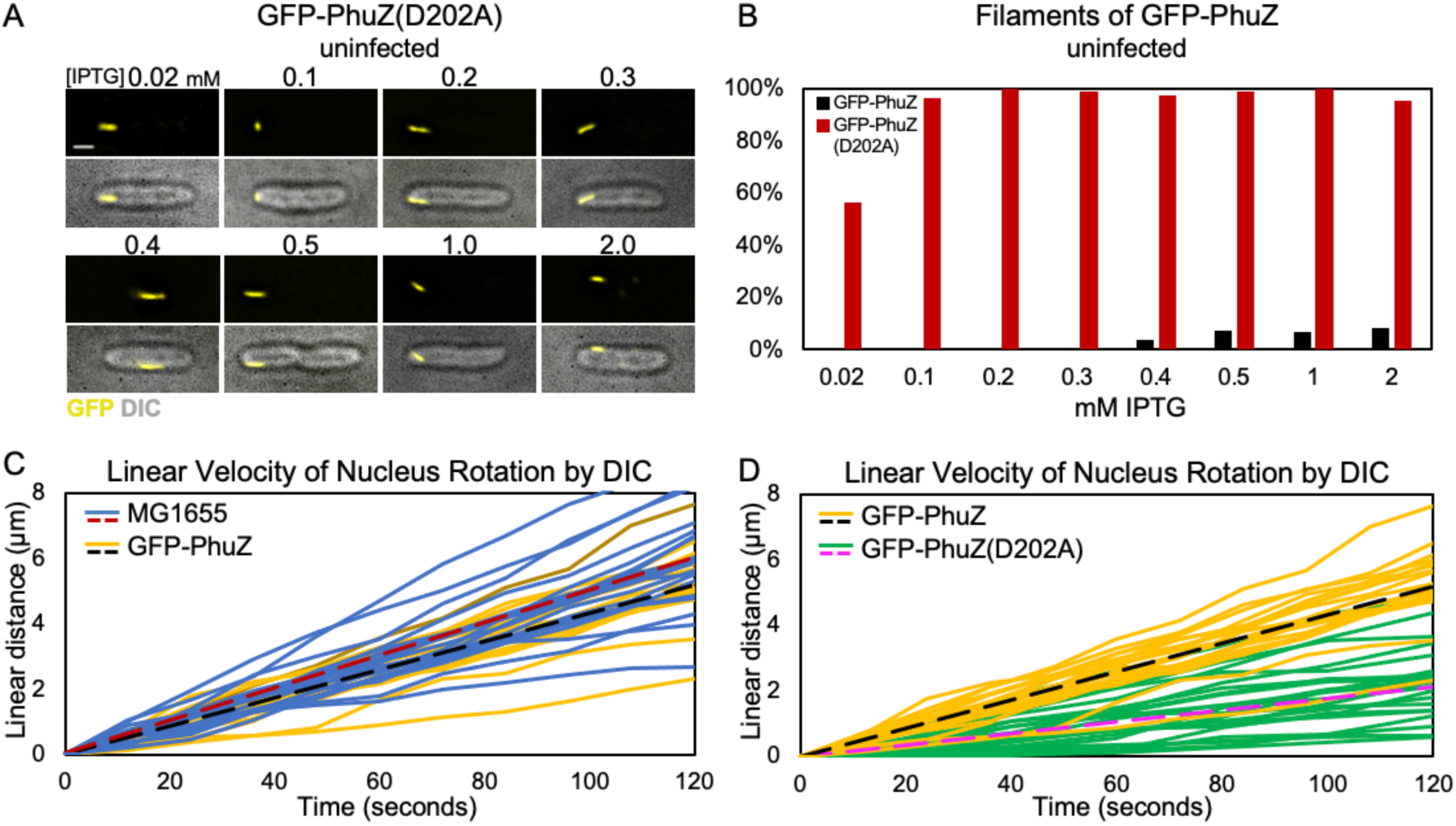
Expression of a catalytically defective PhuZ, GFP-PhuZ(D202A), alters PhuZ assembly properties and phage nucleus rotation. (A) Uninfected *E. coli* (MG1655) expressing GFP-PhuZ(D202A) induced with 0.02-2.0 mM IPTG and grown on 50 μg/ml ampicillin for 90 minutes. Scale bar 1 μm. (B) Percentage of cells expressing either GFP-PhuZ(D202A) (red) or GFP-PhuZ (black) that contained filaments over 0.3 μm at each IPTG concentration (0.2, n = 99; 0.5, n = 85; others, n > 100). (C) Linear velocity of nuclear rotation of 20 measured nuclei for *E. coli* (blue solid lines; average, red dotted line) compared to those in *E. coli* expressing GFP-PhuZ (yellow solid lines; average, black dotted line), 60 minutes after the addition of Goslar. (D) Linear velocity of nuclear rotation of 20 measured nuclei in *E. coli* expressing GFP-PhuZ (yellow solid lines; average, black dotted line) compared to the 23 nuclei in *E. coli* expressing GFP-PhuZ(D202A) (green solid lines; average, magenta dotted line), 60 minutes after the addition of Goslar.

**Figure S5.**
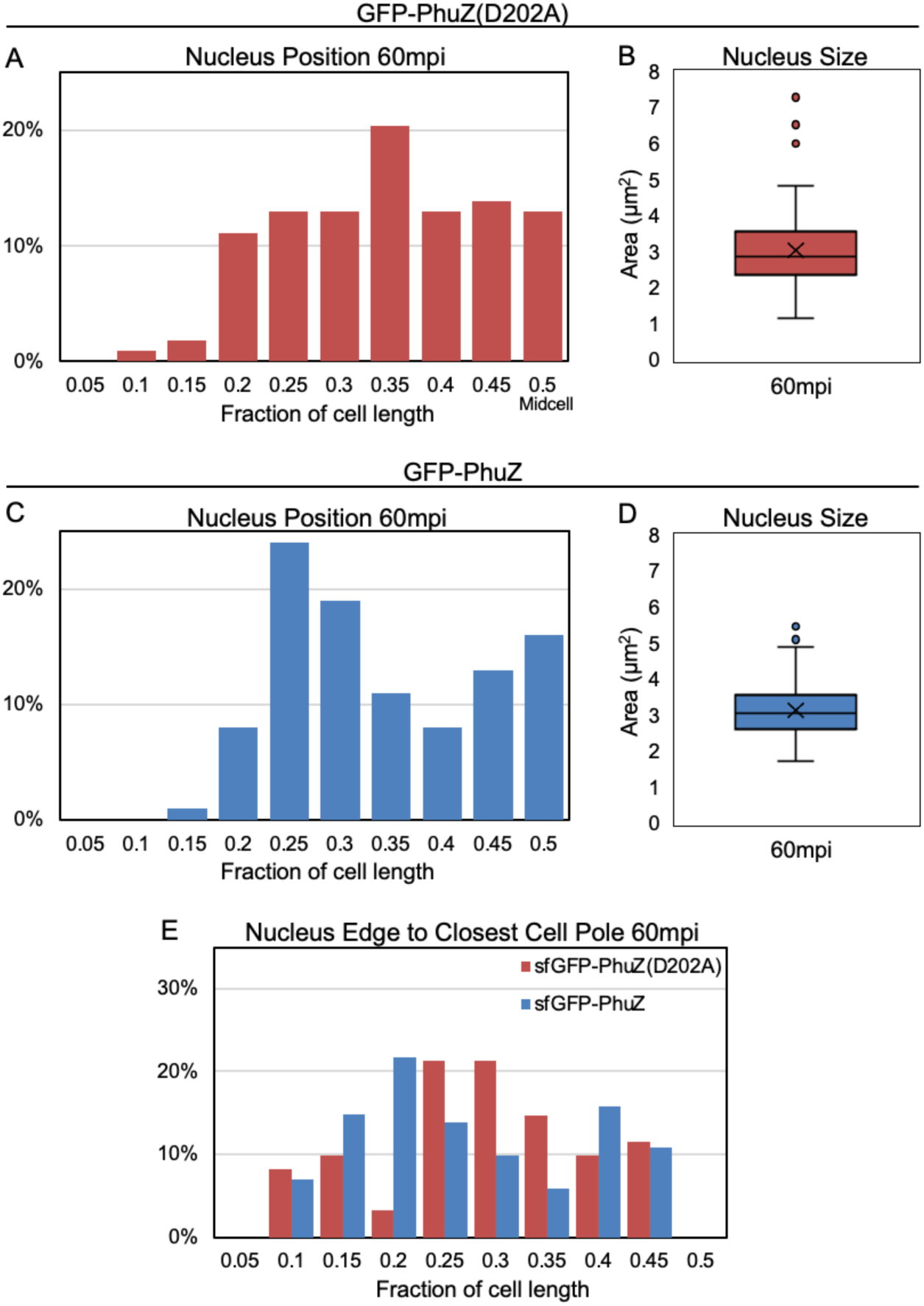
PhuZ(D202A) expression does not affect Goslar nucleus positioning or size. (A) Distribution of 60 mpi phage nuclei (DIC) along the lateral length of *E. coli* (MG1655) expressing GFP-PhuZ(D202A) induced with 0.2 mM IPTG. For each time point, there is no significantly greater chance of finding a phage nucleus in one certain bin than in the neighboring bins (n=108). (B) 2D area of the DAPI-stained 60 mpi phage nucleus in the presence of GFP-PhuZ(D202A) (n=108). (C) Distribution of 60 mpi DIC phage nuclei along the lateral length of *E. coli* expressing GFP-PhuZ induced with 0.2 mM IPTG. For each time point, there is no significantly greater chance of finding a phage nucleus in one certain bin than in the neighboring bins (n=100). (D) 2D area of the DAPI-stained 60 mpi phage nucleus in the presence of GFP-PhuZ (n=111). (E) Distribution of distances between the edge of the phage nucleus to the closest cell pole, normalized to cell length. Either GFP-PhuZ(D202A) or GFP-PhuZ was infected for 60 minutes and stained with FM4-64 and DAPI. DIC was used for measurement (n=88).

**Figure S6.**
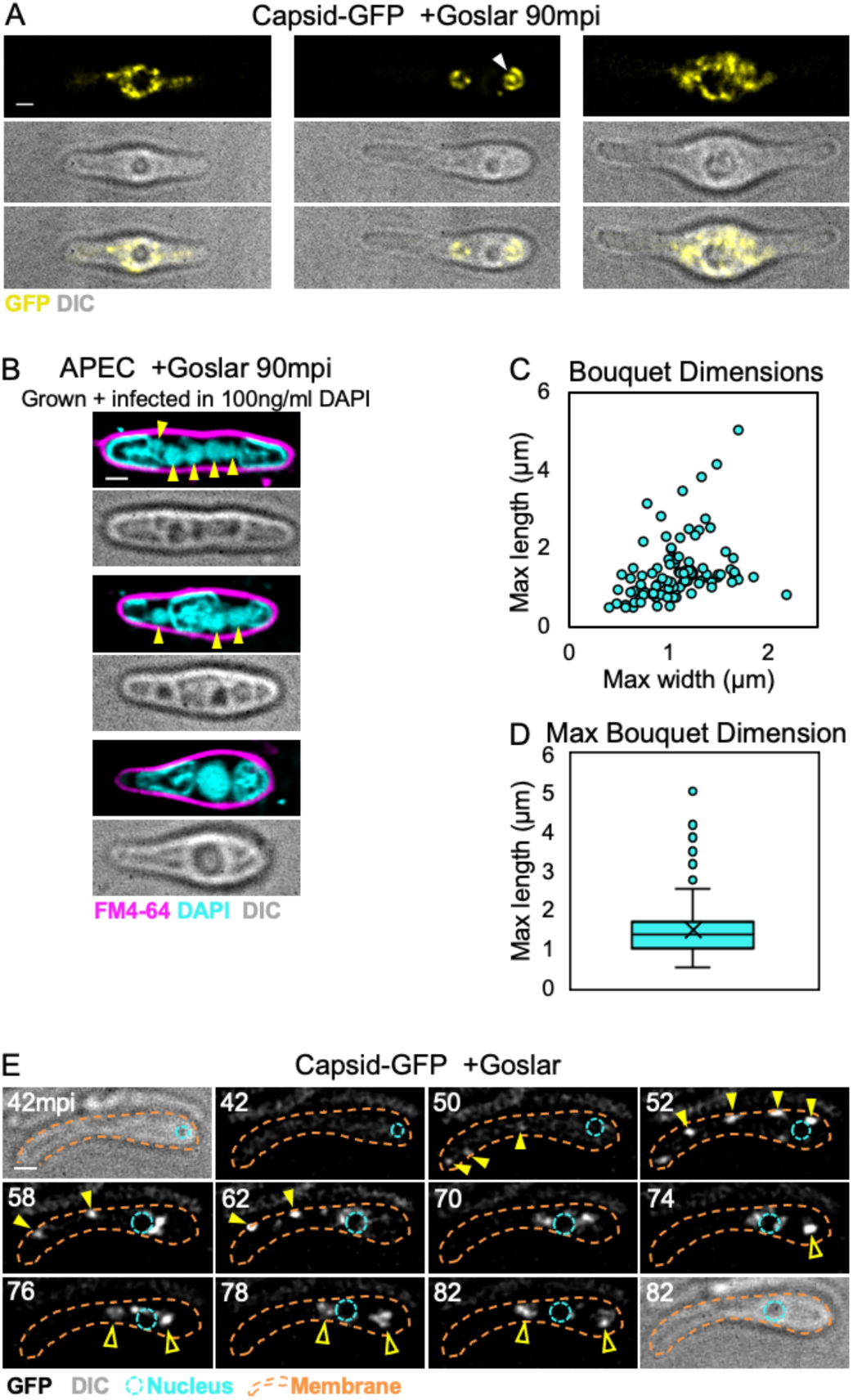
Capsid-GFP localization and bouquet formation during Goslar infections. (A) More examples of capsid-GFP localization as in Figure 7A. *E. coli* (MG1655) expressing the putative capsid protein (gp41) fused to GFP (yellow) induced with 0.2 mM IPTG and infected with Goslar for 90 minutes. Capsids localized around the phage nucleus (left panels), in adjacent bouquets (middle panels), or filling more of the cytoplasm (right panels). All scale bars are 1 μm. (B) Additional examples of multiple nuclei and bouquets (top four panels) and a single nucleus (bottom two panels) imaged by DAPI and FM4-64 after 90 mpi in APEC (yellow arrowheads indicate nuclei). Top 2 panels show 5 nuclei observed in one cell, middle 2 panels show 3 nuclei with a large central bouquet, bottom panels show a single nucleus with the most common bouquet phenotype. (C) DAPI-stained bouquets at 90 mpi in APEC were measured along the maximum x and y dimensions. Each point represents one bouquet (n=105). (D) Box and whisker plot of maximum bouquet dimension for each bouquet in (C). Quartiles are calculated with an excluded median and a mean of 1.5 μm (n=105). (E) Time-lapse of capsid-GFP (white) at various intervals over 40 minutes starting at 42 mpi in MG1655. Capsids migrate from the cell periphery or cytoplasm (yellow solid arrowheads) to the phage nucleus exterior and then to adjacent phage bouquets (yellow open arrowheads) (cyan dashed line, nucleus; orange dashed line, cell membrane).

## Video Legends

**Video 1. GFP-tagged Goslar nucleus rotates in MG1655**.

Live cell GFP time-lapse of a Goslar nucleus at 65 mpi in the strain expressing GFP-shell. Image interval is 2 seconds with a total elapsed time of 30 seconds. Scale bar is 1 μm.

**Video 2. Goslar nucleus rotates without tagged proteins present**.

Live cell DIC time-lapse of MG1655 with no plasmid. Image interval is 2 seconds with a total elapsed time of 60 seconds. Scale bar is 1 μm.

**Videos 3 and 4. Goslar nucleus no longer rotates in the presence of GFP-PhuZ(D202A)**.

Live cell DIC time-lapse of MG1655 expressing GFP-PhuZ(D202A) and infected with Goslar for 75 minutes (Video 3) or 85 minutes (Video 4). Image interval is 4 seconds with a total elapsed time of 80 seconds (Video 3) or 160 seconds (Video 4). Scale bars are 1 μm.

**Videos 5 and 6. Goslar bouquets can contain DNA-filled capsids in the interior**.

3D reconstruction (FIJI – max intensity – interpolated) of DAPI-stained Goslar infections in live cells after 90 mpi. Scale bars are 1 μm.

**Video 7. Goslar capsids migrate from the cell periphery to the nucleus to adjacent bouquets**.

Live cell GFP time-lapse of MG1655 expressing capsid-GFP and infected with Goslar for 10 minutes prior to imaging. DIC images are provided at the beginning and end of the time-lapse for reference of cell boundaries and nucleus position. GFP images begin at 40 mpi with an interval of 2 minutes and a total elapsed time of 40 minutes.

## Notes

### Competing Interest Statement

The authors have declared no competing interest.

